# Complementary assessment of fish biodiversity across the upper/lower mesophotic interface in a subtropical coral reef using eDNA metabarcoding and baited cameras

**DOI:** 10.1101/2025.01.23.634597

**Authors:** Timothy J. Noyes, Leocadio Blanco-Bercial, Stefano Mariani, Gretchen Goodbody-Gringley, Allan D. McDevitt

## Abstract

Mesophotic Coral Ecosystems (MCEs) occur in the middle to lower photic zone (∼30–150 m) of tropical and subtropical regions, are often extensions of shallow reef communities, and generally hold great importance for local commercial fisheries. Compared to their shallower counterpart, MCEs have been traditionally understudied, primarily due to their inaccessibility with traditional monitoring methodologies. In this study, we aim to provide an interdisciplinary assessment of the biodiversity of Bermudan reef fish communities from the upper/lower mesophotic interface (60 m) by utilising a combination of environmental DNA (eDNA) metabarcoding and baited remote underwater videos (BRUVs). In total, 155 species from 137 genera were detected by eDNA metabarcoding whilst a total of 85 species from 53 genera were detected by BRUVs. The combined species detections totalled 182 species with approximately half of those detections unique to this study when compared to previous studies. Both methodologies found differences in α-diversity between study locations, with each method independently detecting the highest species richness at the same location. The species assemblage at each location was dominated (∼80%) by species known to occur throughout the shallow reef system and the upper mesophotic, whilst species only known to inhabit mesophotic ecosystems accounted for ∼6% at each location. These findings suggest a high level of species continuity with the adjacent shallower reef systems. The complementary nature of eDNA metabarcoding and BRUVs allows for a more accurate characterisation of fish biodiversity at the upper/lower mesophotic interface, which can lead to a more comprehensive understanding of ecosystem structure and more informed management decisions.

## Introduction

Providing managers and policy makers with high quality scientific data is key in allowing them to implement effective marine management through an ecosystem-based approach. This is particularly important for Mesophotic Coral Ecosystems (MCEs), which occur in the middle to lower photic zones (∼30–150 m) of tropical and subtropical regions and are often extensions of shallow reef communities (Hinderstein et al., 2010). These reefs harbour high geographic endemism and the emerging consensus is that MCEs have significant ecological importance for reef fishes through providing refuge from fishing pressure and as spawning aggregation sites for shallow reef fish species (Pyle and Copus, 2019).

However, MCEs have been generally understudied in comparison to shallow reef communities, primarily due to inaccessibility with traditional monitoring methodologies (Pyle and Copus, 2019). Environmental DNA (eDNA) is now emerging as a proven and valuable tool for ecosystem monitoring in a wide range of environments (Deiner et al., 2017; Taberlet et al., 2018) including marine ecosystems (Miya et al., 2020; Mathon et al., 2022). However, government agencies are often not willing to change monitoring methodologies before proof of concept has been established. Environmental DNA has the potential to provide a more “complete” biodiversity assessment than other methods due to its efficiency in detecting cryptic, low-abundance, transient and rare taxa (Boussarie et al., 2018; Gold et al., 2021). In addition, the economic efficiency of eDNA metabarcoding allows biomonitoring to be conducted at large spatial and temporal resolutions which when paired with biotic measurements, would greatly benefit ecosystem-based management approaches (Yao et al. 2022). Despite the benefits of the eDNA metabarcoding approach, there are still uncertainties and artefacts inherent to the methodology. The samples often contain a combination of degraded target taxa DNA (Collins et al. 2018) and non-target DNA that may co-amplify (Stat et al. 2017); inhibition of DNA amplification or the introduction of false positives throughout multiple stages (e.g., sampling, DNA extraction, amplification, and sequencing) can lead to the misinterpretation of sequencing data sets; all of which still hinders the systematic application of eDNA metabarcoding for routine monitoring.

Baited Remote Underwater Video systems (BRUVs) are non-destructive, cost-effective fishery-independent sampling units (Langlois et al., 2010) that produce spatially explicit quantitative data on reef ichthyofauna abundance and diversity. BRUVs techniques can survey a broad range of species (increased through the presence of bait; Harvey et al., 2007; Dorman et al., 2012) and as such have been utilized in various marine environments (Whitmarsh et al. 2017), including for the assessment of pelagic species (Santana-Garcon et al. 2014). In addition, the video footage creates a permanent record that is especially useful for life stage and behavioural observations (Barley et al. 2016), allowing third-party verification of species identification and for use in education and outreach initiatives. Whilst the presence of bait can increase species richness, it can also be biased towards certain trophic guilds (i.e., carnivores; Stobart et al. 2007) leading to underrepresentation of smaller cryptic species (Lowry et al. 2012). Screen saturation (Schobernd et al. 2014) due to abundant taxa can also lead to an underestimation of the population due to a physical number of individuals only be able to fit in the field of view. In addition, it is not always possible to distinguish between closely related species.

Bermuda’s coral reef system transitions through a series of shallow patch reefs, rim reefs, and terraced reefs (Logan 1988; Logan and Murdoch 2011) dropping quickly to deeper mesophotic reefs that surround the main reef platform (Goodbody-Gringley et al., 2019a). With the exception of Stefanoudis et al. (2019), previous Bermudan mesophotic fish community investigations have been single methodology approaches (Pinheiro et al. 2016) and primarily focused on invasive species assessments (Goodbody-Gringley et al. 2019a, 2023). The valuable ecological services (Holmlund and Hammer 1999) that MCEs provide in Bermuda are under threat due to the presence of two invasive lionfish species, red lionfish (*Pterois volitans*) and devil firefish (*P. miles*) that aggregate on MCEs (Goodbody-Gringley et al. 2023). Studies have reported that greater lionfish abundances correlate with significant declines in prey fishes (Albins and Hixon 2008; Morris and Green 2012).

This study provides an interdisciplinary assessment of fish biodiversity in Bermuda’s upper/lower mesophotic interface (∼60–65 m) by utilising a combination of eDNA metabarcoding and BRUVs (Stat et al., 2019; Aglieri et al., 2021). This depth zone has previously been quantified as a faunal break in other mesophotic regions (e.g., Pinheiro et al., 2016; Pyle et al., 2016; Lesser et al., 2019) and, locally, it remains an important area for local commercial fisheries, such as spiny lobsters (*Panulirus argus*) and both demersal and pelagic fin fishes (Faiella, 2003). We aim to study trends in the wider Bermudan upper/lower mesophotic interface by investigating spatial and temporal variation in α- and β-diversity patterns across both survey methods.

## Materials and methods

Surveys were conducted monthly between August – December 2017, and every two months between May and October 2018. Site locations followed the 60 m depth contour and were situated ∼350 m apart (Fig. 1) to maintain site fidelity during BRUVs deployments by minimising the influence of bait plumes between sites (Harvey et al. 2007). Active steps to minimise cross contamination between eDNA sampling and BRUVs bait were taken at all times through the following procedures: (a) seawater samples were collected prior to BRUVs deployments; (b) assisting personnel were each assigned to only one methodology; (c) external surfaces of eDNA storage coolers and BRUVs equipment were sprayed with 10% bleach solution and rinsed prior to loading on sampling vessel and post deployments; (d) sampling equipment was kept in separate locations on the sampling vessel and (e) BRUVs bait was kept frozen in a sealed container until use.

**Fig. 1.**
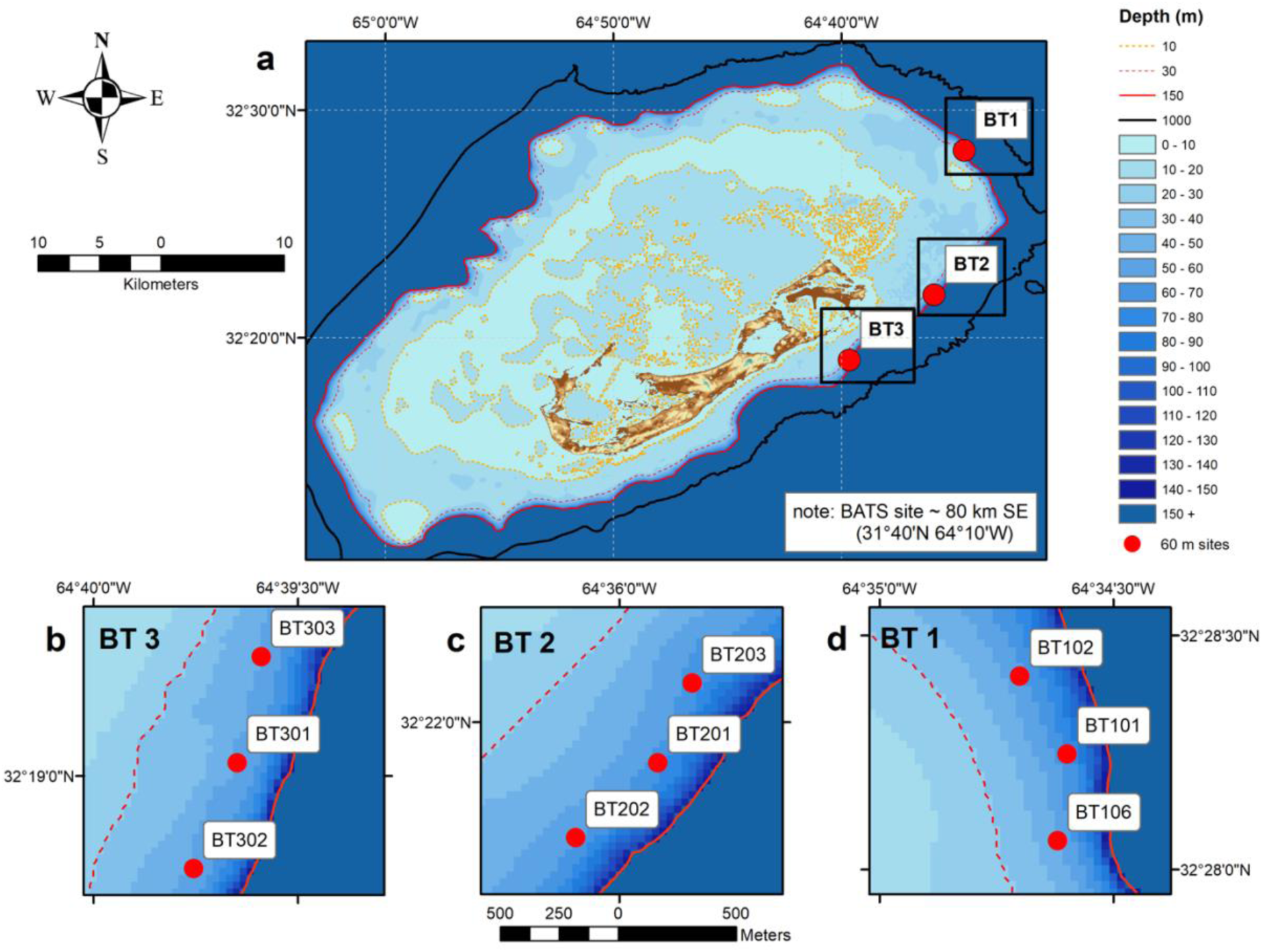
(**a – d**), 30 m depth contour dashed red line, 150 m depth solid red line, (**a**). Map of Bermuda and study locations (red circles), 10 m contour orange dotted line, 1000 m contour solid black line. (**b**) Bathymetric map of Location BT3, red circles = 60 m sites, (**c**) Bathymetric map of Location BT2, red circles = 60 m sites, (**d**) Bathymetric map of Location BT1, red circles = 60 m sites

### Environmental DNA collection

In brief, sample seawater was collected using a 12 L Niskin lowered to a target depth (∼ 2 m above the benthos) with site depth verified via boat-mounted depth sounder. All sampling and filtering equipment were rinsed three times with 10% bleach solution and then three times with milliQ (Miya et al. 2015) prior to the collection of samples. A minimum volume of 8 L was collected to reduce false negative detection probabilities (Yao et al. 2022). Sample seawater was drawn into 2 x 4L Nalgene carboys that were flushed three times with sample seawater prior to filling. All sample carboys were stored on ice inside black bin bags within coolers until back at the laboratory, which ranged from 5 – 6 hours depending on the distance of the sampling location from the laboratory facilities. Despite a recent study by Mächler et al. (2018) finding no discernible effect of sunlight or UV on the detectability of eDNA, we took the precaution of cooling the samples and limiting exposure to UVB radiation to reduce DNA degradation (Strickler et al. 2015). In the interest of maintaining a standardised procedure, the practice of keeping sample bottles in bags on ice out of direct heat and light was the standard operating procedure.

At the first available opportunity upon return to the laboratory facilities, samples were filtered through inline filter holders loaded with 47 mm 0.8 μm hydrophilic membrane polycarbonate (Takahara et al. 2012) filters (NucleporeTM Track-Etched Membrane (Whatman®)) using a vacuum filtration set up. Sample carboys remained housed within the coolers (i.e., in the dark on ice) until the filtration process for a specific carboy had begun. On average, filtration took 3 hours per 4 L of sample. Ideally, the full sample volume (8 L) was passed through a single filter, however during the high productive months (higher particle load due to increased primary productivity; May – August) an additional filter was required to achieve the desired filtered sample volume (8 L) due to the retention of additional suspended matter (phytoplankton, detritus, etc.) clogging the filters (Goldberg et al. 2016). Filters were individually stored in 3.6 ml cryogenic vials (USA Scientific, FL, United States) together with 1 mL of sterile 3 mm porcelain spheres (Rio Grande, CA, United States), and stored at −23°C in a dedicated eDNA freezer until extraction. All filtration activities were conducted in laboratory space designated for seawater filtration that had been presterilized (10% bleach wash) to minimise contamination.

### Laboratory methods

To minimize contamination, separate laboratories were used for eDNA extractions, pre-PCR and post-PCR procedures, with all laboratory spaces being cleaned prior to and post sample processing. DNA extractions from the frozen filters were performed using the E.Z.N.A. Mollusc DNA Kit (Omega Bio-Tek: Norcross, Georgia, USA). See the Supplementary Material for details on modifications to the protocol. To confirm successful DNA extraction, total DNA concentration (i.e., fish plus non-target DNA) and purity of each sample were quantified using a NanoDrop One Microvolume UV-Vis Spectrophotometer (Thermo Scientific: Waltham, Massachusetts, USA).

Library preparation was performed as a two-step Polymerase chain reaction (PCR) using AccuPower ProFi Taq PCR PreMix (Bioneer: California, USA) and the MiFish-U teleost-specific primers (12S ribosomal RNA gene 163-185 bp; Miya et al., 2015). See the Supplementary Material for full details of the PCR and library preparation protocols. In total, 67 biological and 31 control (field (n = 6), extraction (n = 5) and PCR blanks (n = 13), and positive controls (n = 7)) samples were sequenced.

Sequencing protocols were as follows: library concentration was determined using Qubit Fluorometer (Thermo Fisher Scientific) and quality assessed with an Agilent Fragment Analyser (Agilent, Santa Clara, CA). The library was normalized to equimolar concentrations alongside another one from a separate, concomitant study, and both libraries were then pooled for size selection (200 – 400 bp) using a Pippin HT (Sage Science). To normalise pooled libraries, concentrations were confirmed, and quality assessed using a Qubit Fluorometer (Thermo Fisher Scientific) and an Agilent Fragment analyser (Agilent, Santa Clara, CA). Sequencing was performed using a V2 Reagent Kit and standard flow cell with paired end reads of 250 bp with custom sequencing primers on an Illumina MiSeq platform (Illumina, San Diego, CA, USA).

### Bioinformatics

Illumina’s Binary Base Call (BCL) Convert software was used to convert the raw BCL files generated by the sequencing process to FASTQ files and separate individual samples based on dual index barcodes. Demultiplexed samples were analysed in MOTHUR version 1.44.3 (Schloss et al. 2009) following an updated pipeline (Blanco-Bercial 2020). Contigs were aligned with a Phred quality score threshold of >30 and any resulting reads that were shorter than 115 bp or had ambiguities were removed. Retained unique sequences were aligned against the 12S region from the MitoFish Reference Database (Sato et al. 2018) and trimmed to the length of the region. All incomplete reads, those that did not reach both ends of the alignment were discarded. Chimeric sequences were removed using VSEARCH (Rognes et al. 2016) implemented within MOTHUR. Single variants (=100% Molecular Operational Taxonomic Units;Porter and Hajibabaei, 2018) were obtained using DEBLUR implemented within MOTHUR (differences =1). The resulting variants were taxonomically identified by BLASTing again the GenBank nucleotide database. Taxa similarity thresholds were set at 99% for species level assignments (Stat et al. 2019; Juhel et al. 2020) with the exception of *Abudefduf* (Forsskål 1775), *Mulloidichthys* (Whitley 1929) and *Thalassoma* (Swainson 1839) that were set at 98% due to only a single species from each genus having been verified and documented in Bermuda to date. Retained MOTU assignments were passed through a second step of taxonomic confirmation through comparison to a custom-made database of fishes native to Bermuda’s marine environment (Noyes and Blanco-Bercial unpublished; see appendices for details on FASTA sequences). BLAST results were manually checked, and taxonomic nomenclature was based on the World Register of Marine Species (WoRMS; http://www.marinespecies.org/).

A contamination threshold approach-based was applied to the raw MOTU community data (Deiner et al. 2017). The average read count of MOTUs detected in controls samples was used as the correction factor specifically for that species (Bokulich et al. 2013; Port et al. 2016; Deiner et al. 2017) and applied to all environmental samples. That is, taxa with read counts below correction factor values were removed from corresponding environmental samples. All non-fish species were removed prior to final analyses.

### Baited Remote Underwater Video systems (BRUVs)

Single GoPro™ Hero 3+ cameras in Golem Gear underwater housings (1000 m depth rated) were mounted inside purpose built galvanized steel frames. The frames provided multiple functions by allowing cameras to be mounted at a set distance (45 cm) from the benthos, acting as ballast during deployments, and providing protection for the camera housings during deployment and recovery. All cameras were set to record at 1080 definition on medium field of view with each system baited with ∼ 700 g of chopped redear herring, (*Harengula humeralis*) placed in plastic mesh and suspended ∼ 1.5 m in front of the cameras. The bait bag was suspending on flexible conduit to allow the bait to maintain contact with the benthos. Having the bait in contact with the benthos has been known to increase sightings of crypto-benthic species (M. Cappo; pers. comm.). Systems were left to record on the seafloor for a minimum of 1 hour between the hours of 11:00 – 16:00. All recordings were made using ambient light. Video footage was annotated using EventMeasure software (www.seagis.com.au) which is specifically designed for logging and reporting events that occur in digital imagery. The measurement matrix generated during the annotations was relative abundance and defined as the maximum number of individuals per species seen at once during a 60-minute video or “MaxN”. The 60-minute observation period began once the system had become stable after contacting the benthos with fish species identified to the lowest taxonomic level possible. The exception to this level of identification was for lionfish (*Pterois* spp.). Whilst the dominant species in Bermuda is the red lionfish (*P. volitans*), it is not possible to visually distinguish between its congener the devil firefish (*P. miles*) with 100% certainty. Therefore, they have both been pooled to genus level.

It should be noted that any bait-related biases such as the area of attraction or species attracted are deemed constant throughout the dataset. Whilst every attempt was made to survey the same sites through deployment at the same latitude and longitude (Garmin GPS +/-5 m accuracy), the orientation of BRUVs cannot be controlled for during deployment and is therefore of a stochastic nature (Fig. 2). Habitat metrics were visually estimated from BRUVs imagery for each deployment and composed of slope, relief (structural complexity) and nine benthic categories (rubble and sand, hard bottom, live coral, gorgonians, crustose coraline algae (CCA), rhodoliths, macroalgae, turf algae, sponges). Benthic categories were estimated as percentage cover, relief was categorised as per the six-point scale of Wilson et al. (2007) and slope was graded on a six-point scale from flat to vertical. It was deemed not appropriate to assign set habitat metrics to a site due to a lack of knowledge on Bermudan mesophotic benthic community seasonality and the stochastic nature of deploying BRUVs to 60 m. Abiotic variables generated in Noyes et al., (2024) were matched to sampling sites used for this assessment (Table 1). See Noyes et al., (2024) for explicit details on how these abiotic data were derived.

**Fig. 2.**
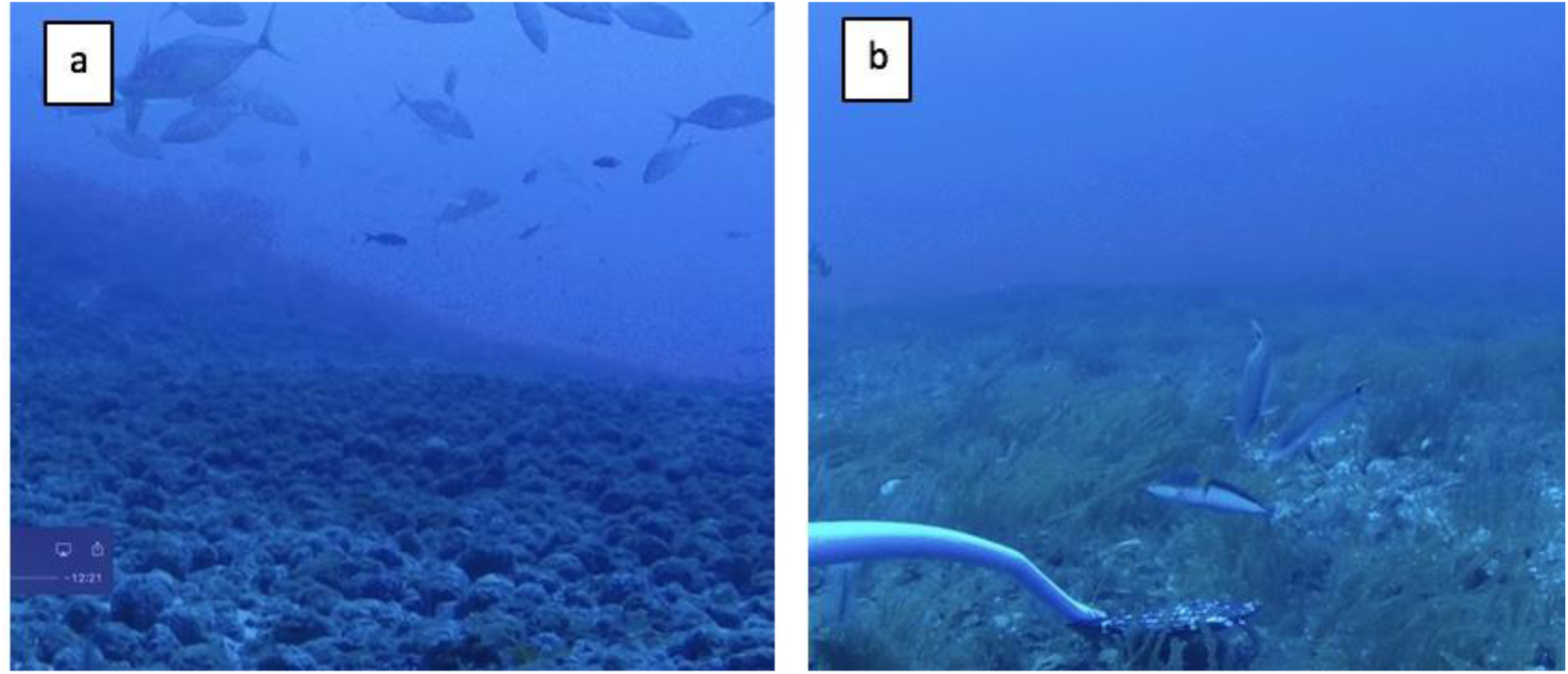
Seabed complexity observed at site BT1 on survey 1 (**a**) and survey 4 (**b**) differences are caused by changes in BRUVs orientation upon contact with the seabed (depth ∼ 60 m). Image (**a**) depicts rhodolith beds foreground with ∼ 5 m reef structure in the background and a shoal of Gwelly jacks (*Pseudocaranx dentex*) in the field of view. Image (**b**) depicts rhodolith beds and brown algae (*Sporochnus bolleanus*), two Sand tilefish (*Malacanthus plumieri*) and one Yellowhead wrasse (*Halichoeres garnoti*).

**Table 1.** Summary of abiotic and biotic explanatory variables used for Redundancy Analysis (RDA)

| Abbreviation | Description | Abbreviation | Description |
| --- | --- | --- | --- |
| Ar | $\Omega$ Aragonite | CCA | Crustose coralline algae |
| Ca | $\Omega$ Calcite | coral | Living coral |
| DIC | Dissolved Inorganic Carbon | gorgonians | Gorgonian |
| NAO | North Atlantic Oscillation index | hard bottom | Hard bottom |
| NEC | Net Ecosystem Calcification | macroalgae | Macroalgae |
| NEP | Net Ecosystem Productivity | rhodolith | Rhodolith |
| pH | pH | sediment | Unconsolidated benthos |
| PSU | Practical Salinity Units | turf | Turf algae |
| Revelle | Revelle Factor | Sum 17 | Summer 2017 |
| T | measured temperature (°C) at time of eDNA collection | Win 17 | Winter 2017 |
| TA | Total Alkalinity | Spr 18 | Spring 2018 |
| Slope | Reef slope | Sum 18 | Summer 2018 |
| relief | relief | Aut 18 | Autumn 2018 |

### Data analysis

All downstream analyses were performed in R version 3.5.3 (R Core Team 2019) using the packages adespatial v0.3-7 (Dray et al. 2021), betapart v1.5.4 (Baselga and Orme 2012), FSA (Ogle et al. 2021), iNEXT_2.0.20 (Hsieh et al. 2016) and vegan v2.5-6 (Oksanen et al. 2019). To be consistent with the derived taxonomic assignments from the BRUVs, eDNA detections of both lionfish spp. (*P. volitans* and *P. miles*) were pooled to genus level.

Variation in α- (taxon richness at one site) and β- (changes in taxon composition) diversity were investigated between assessment methodologies (eDNA, BRUVS) whilst β-diversity was also considered at the genus level.

Kruskal-Wallis and Dunn’s tests (p-values adjusted by the Benjamini-Hochberg Procedure) from the package’s stats v3.5.3 and FSA v0.9.1 (Ogle et al. 2021) respectively were used to compare α-diversity between location (BT1, BT2, BT3) and between season (summer 2017, autumn 2017, winter 2017, spring 2018, summer 2018, autumn 2018) for both assessment methodologies due to nonnormal distribution of data (Shapiro–Wilk normality test: W = 0.903, P = < 0.001). The iNEXT package was used to compute sample size-based rarefaction and extrapolation curves using incidence frequencies of taxa for 50 samples, 10 knots, 1000 bootstraps and 95% confidence intervals.

Beta diversity (using Jaccard dissimilarity) was separated by sample methodology and partitioned into taxon replacement (turnover) or taxon subgroups (nestedness-resultant). Community heterogeneity for the respective β-diversity components were grouped by location and compared by calculating homogeneity of multivariate dispersions (MVDISP) and statistically tested using ANOVA.

Dissimilarity matrices for β-diversity components (Jaccard dissimilarity) partitioned by sample methodology were created to test the effects of location, site (BT101, BT102, BT106, BT201, BT202, BT203, BT301, BT302, BT303, Fig 1 b-d) and season, and were tested statistically using permutational multivariate analysis of variance (PERMANOVA). The total β-diversity component for both sample methodologies were graphically represented using non-metric MultiDimensional Scaling (nMDS).

To determine if species detections for both methodologies were influenced by environmental forcings (explanatory variables of benthic community composition, abiotic conditions, and seasonality), a canonical form of principal component analysis (redundancy analysis; RDA) was applied to each dataset independently and visualized with a triplot (RDA biplots with explanatory variables plotted as arrows). Both a global test and test of the canonical axes with 999 permutations were performed on the resultant RDA outputs. Prior to the implementation of the RDA, a Hellinger transformation was applied to both the BRUVs and eDNA species datasets whereby abundance values are divided by the sample total abundance and then square-root transformed (Legendre and Gallagher 2001). The environmental variables were standardised to zero mean and unit variance (Table 1).

## Results

In total 67 paired collections of seawater for eDNA analysis and BRUVs deployments met the sampling criteria and were included in the downstream analyses. The 12S amplicon libraries produced 6,360,726 raw reads, available from BioProject PRJNA1205382. After the removal of singletons, the application of the minimum read count threshold (5 reads) and assignment to genus level, the number of reads was reduced to 2,097,857. When taxa assignments were further refined to species level, reads where reduced to 769,623 in total. Overall, there were a total of 155 species from 137 genera detected by eDNA metabarcoding (Fig. 3; Table S1). Importantly, the bait utilised for the BRUVs was not detected (redear herring *Harengula humeralis*) by eDNA. Baited cameras detected a total of 85 species from 53 genera which equated to ∼45% less species and ∼60% less genera than eDNA (Table S1). The greater detection rates of fish taxa by eDNA were consistent at both the species and genus level across all three survey locations (Fig. 3). Sample size-based rarefaction/extrapolation (R/E) curves (Fig. S1) imply taxon diversity had reached a plateau at all locations which would infer a suitable level of sampling had been reached to assess the community structure of these three mesophotic locations for both methodologies.

**Fig. 3.**
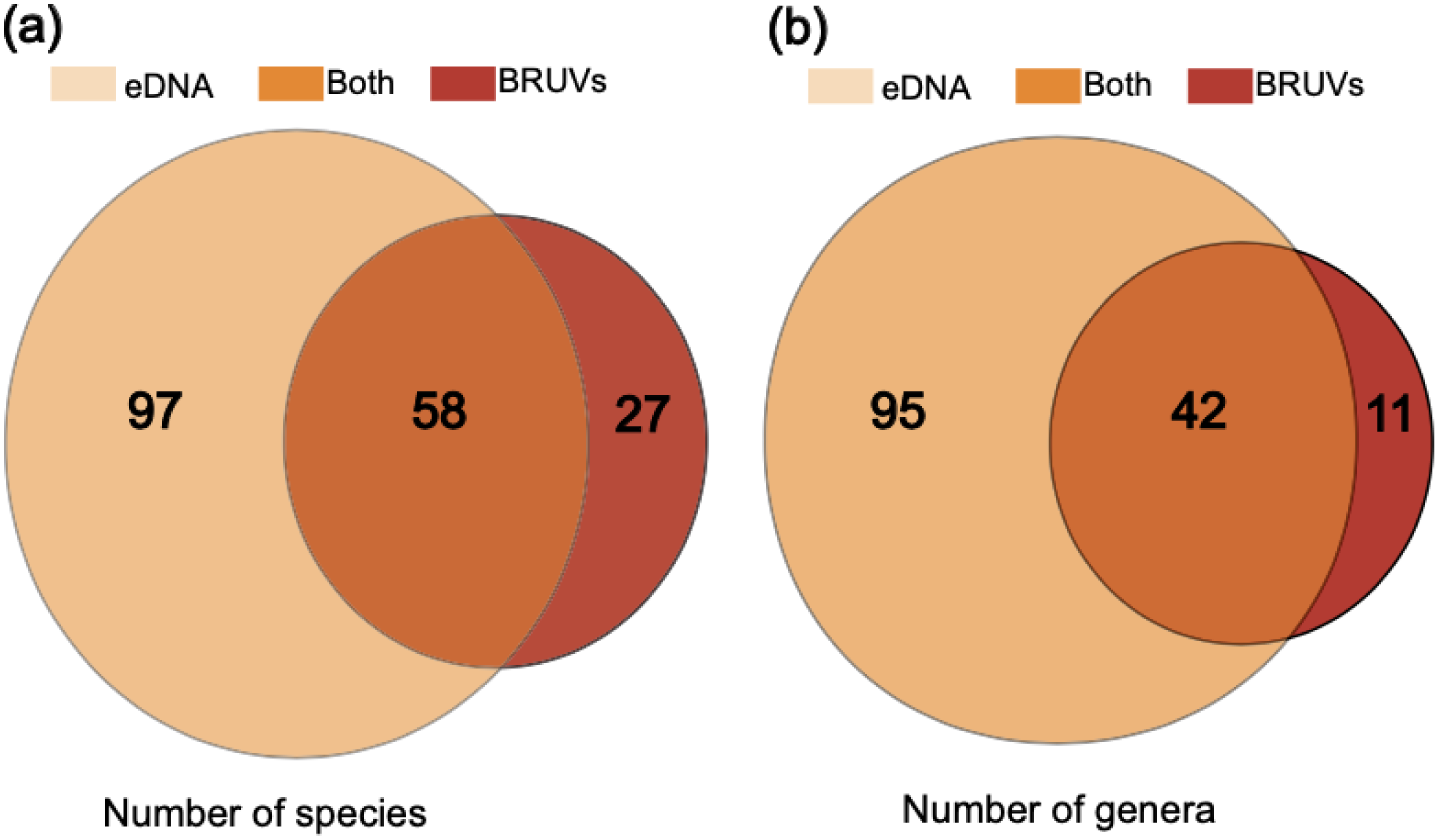
Venn diagrams of the number of species (**a**) and genera (**b**) detected by the two assessment methodologies, Baited Under Water Video systems (BRUVs; red ellipses) and environmental DNA (eDNA; peach ellipses). The orange intersect indicates the number of species (a) and genera (b) detected by both methodologies.

Differences in α-diversity between locations were detected by both eDNA (Kruskal-Wallis test, H2 = 12.582, P = 0.002) and BRUVs (Kruskal-Wallis test, H2 = 6.423, P = 0.040; Fig. 4a) with lower taxon richness measured at BT2 than BT3 (Dunn’s test, Z = −3.527, P.adj = 0.001) by eDNA and lower taxon richness measured by BRUVs at BT1 than BT2 (Dunn’s test, Z = −2.421, P.adj = 0.046). The remaining between-location taxon richness comparisons were comparable. Within location taxon differences were only recorded by BRUVs and only within the BT3 location. This was due to lower α-diversity at site BT302 (n = 27; Dunn’s test, Z = 3.233, P.adj = 0.004) than BT302 (n = 50) and site BT303 (n = 43; Dunn’s test, Z = −3.041, P.adj = 0.004) respectively.

**Fig. 4.**
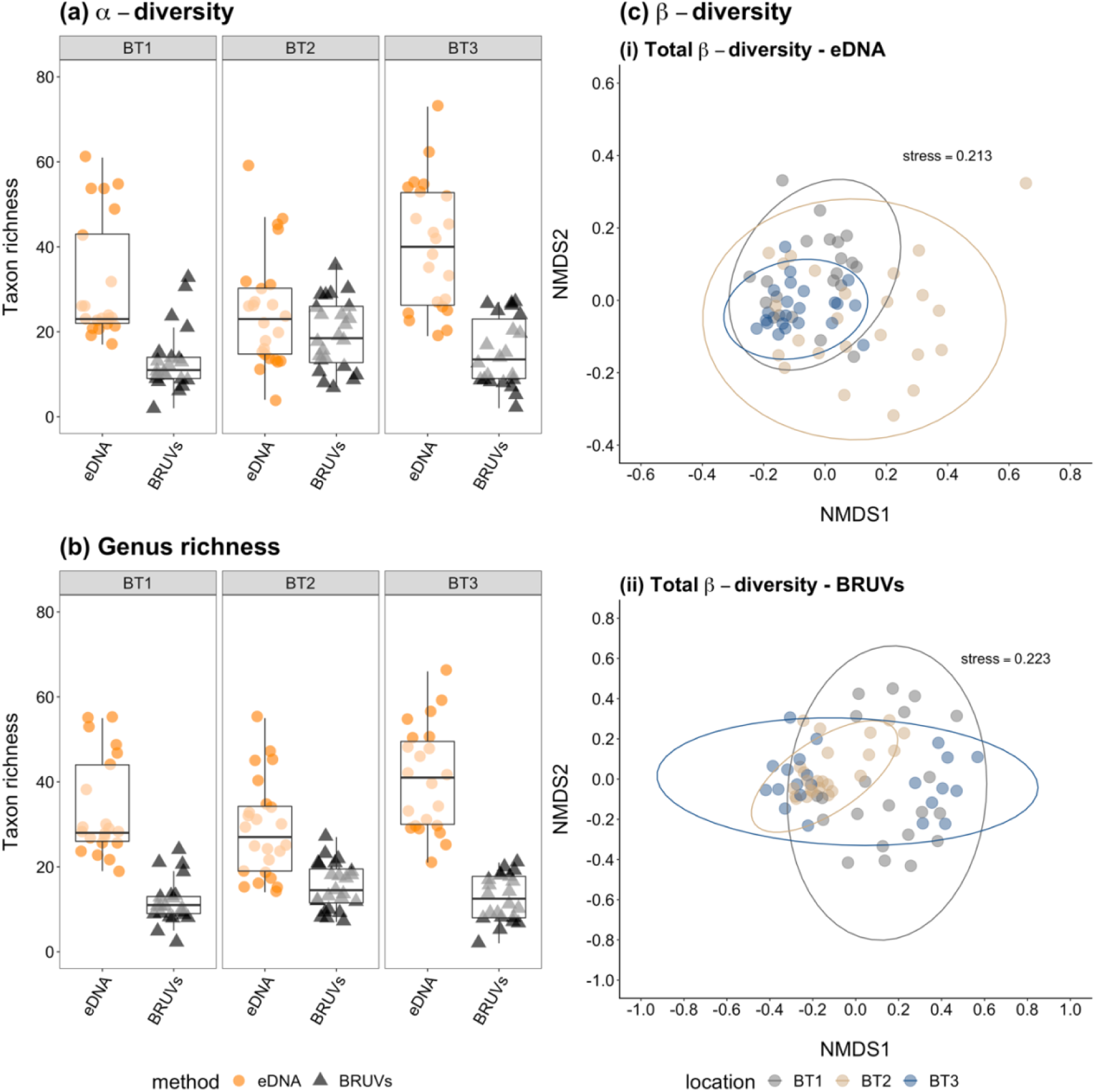
Summaries of α- and β-diversity between locations partitioned by sampling methodology: (**a**) boxplot of species richness, boxes indicating 25th, 50th and 75th percentiles and whiskers show 5th and 95th percentiles, eDNA (orange circles) and BRUVs (dark grey triangles). (**b**) boxplot of genus richness, boxes indicating 25th, 50th and 75th percentiles and whiskers show 5th and 95th percentiles, eDNA (orange circles) and BRUVs (dark grey triangles) (**c**) non-metric multidimensional scaling plots of β-diversity components, (i) eDNA and (ii) BRUVs, locations: BT1 (dark grey symbols and ellipses), BT2 (peril symbols and ellipses), BT3 (blue symbols and ellipses).

Seasonal differences in α-diversity were detected by eDNA at locations BT2 (H4 = 14.37, P = 0.006) and BT3 (H5 = 15.485, P = 0.008) but not BT1 (H5 = 10.683, P = 0.059). Higher taxon richness was detected in spring (2018: Dunn’s test, Z = 2.545, P.adj = 0.036) than summer at BT2; at the same location, taxon richness was higher in summer 2017 than summer 2018 (Dunn’s test, Z = 2.821, P.adj = 0.048). At BT3, autumn 2017 taxon richness was greater than the summer 2018 (Dunn’s test, Z = 2.486, P.adj = 0.048).

The degree of variation in species composition (i.e., total β diversity) between locations were significantly different for both sampling methodologies. Beta diversity at each of the three locations was primarily driven by species replacement and not species loss. The percentage of species turnover across each location were comparable for both datasets.

With the exception of species loss (nestedness-resultant) within the BRUVs dataset, the interaction of location did have a significant influence on all components of β diversity for both datasets (Table 2). The influence of site was only detectable within the BRUVs dataset (Table 3) with a separation in ordination between sites within BT1 and BT3 samples within BT3 (Figure 4 cii). The influence of seasonality was only significant on eDNA derived community data (Table 3).

**Table 2.**
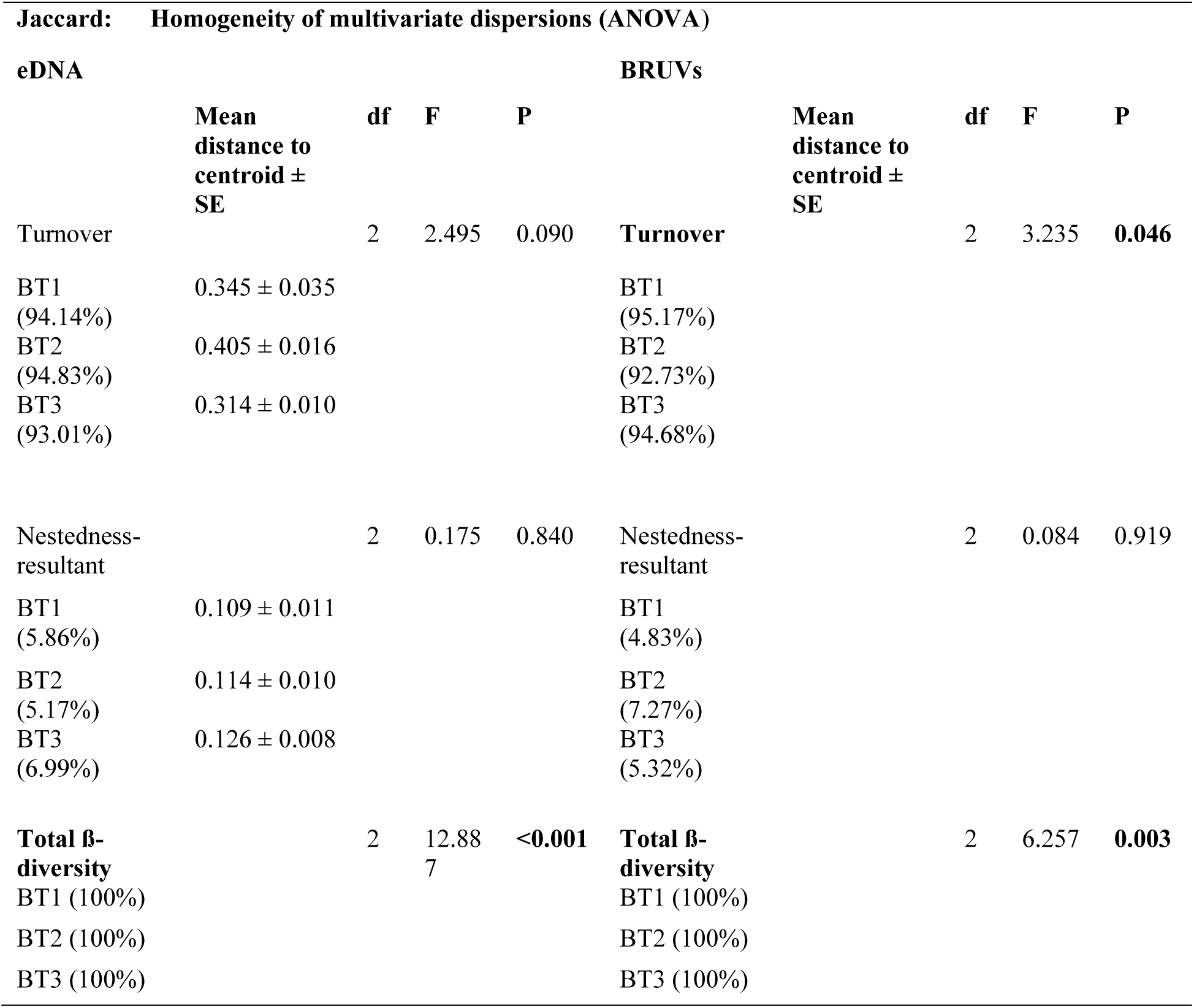
Summary statistics of community heterogeneity for β-diversity (Jaccard dissimilarity) components (turnover, nestedness-resultant) partitioned by sampling methodology (eDNA metabarcoding, BRUVs) grouped by location (BT1, BT2, BT3) statistically tested using ANOVA. Statistically significant results are indicated in bold text. The relative contributions of the respective β-diversity components for each location are given in brackets.

**Table 3.** Summary statistics of the interactions of location (BT1, BT2, BT3), site (BT101, BT102, BT106, BT201, BT202, BT203, BT301, BT302, BT303) and season (summer 2017, autumn 2017, winter 2017, spring 2018, summer 2018, autumn 2018) on community composition partitioned by β-diversity components (Jaccard dissimilarity) separated by sampling methodology (eDNA metabarcoding, BRUVs) statistically tested using permutational multivariate analysis of variance (PERMANOVA). Statistically significant interaction results are indicated in bold text.

|  |  | Jaccard: |  | Community similarity (PERMANOVA) |  |  |  |  |  |
| --- | --- | --- | --- | --- | --- | --- | --- | --- | --- |
| eDNA |  |  |  |  | BRUVs |  |  |  |  |
| Interaction | df | F | R<sup>2</sup> | P | Interaction | df | F | R<sup>2</sup> | P |
| Turnover |  |  |  |  | Turnover |  |  |  |  |
| location | 2 | 1.994 | 0.064 | **0.005** | location | 2 | 6.702 | 0.166 | **0.001** |
| site | 6 | 0.883 | 0.084 | 0.713 | site | 6 | 2.782 | 0.206 | **0.001** |
| season | 5 | 1.309 | 0.104 | 0.092 | season | 5 | 1.042 | 0.064 | 0.411 |
| location:season | 9 | 0.657 | 0.094 | 0.979 | location:season | 9 | 1.087 | 0.121 | 0.360 |
| site:season | 25 | 0.880 | 0.351 | 0.788 | site:season | 25 | 0.675 | 0.208 | 0.988 |
| Nestedness-resultant |  |  |  |  | Nestedness-resultant |  |  |  |  |
| location | 2 | 6.155 | 0.096 | **0.014** | location | 2 | -<br>6.150 | -0.140 | 0.991 |
| site | 6 | 0.779 | 0.036 | 0.602 | site | 6 | 2.224 | 0.152 | 0.267 |
| season | 5 | 6.681 | 0.261 | **0.001** | season | 5 | 3.615 | 0.206 | 0.153 |
| location:season | 9 | 3.628 | 0.255 | **0.014** | location:season | 9 | -<br>0.181 | -0.019 | 0.917 |
| site:season | 25 | 1.040 | 0.203 | 0.485 | site:season | 25 | 2.053 | 0.584 | 0.240 |
| Total β-diversity |  |  |  |  | Total β-diversity |  |  |  |  |
| location | 2 | 2.811 | 0.078 | **0.001** | location | 2 | 4.296 | 0.115 | **0.001** |
| site | 6 | 0.859 | 0.071 | 0.902 | site | 6 | 2.241 | 0.181 | **0.001** |
| season | 5 | 2.249 | 0.155 | **0.001** | season | 5 | 1.036 | 0.070 | 0.363 |
| location:season | 9 | 1.098 | 0.137 | 0.193 | location:season | 9 | 0.978 | 0.118 | 0.541 |
| site:season | 25 | 0.859 | 0.297 | 0.975 | site:season | 25 | 0.778 | 0.261 | 0.997 |

The factors included in the redundancy analysis accounted for approximately half of the variance within each fish community matrix (eDNA; R^2^ = 0.502, BRUVs; R^2^ = 0.495). Overall, both assessments showed there to be a degree of separation between the three locations, however this was most evident in the BRUVs derived dataset (Fig. 5). The associations between the environmental metrics and the communities were not consistent between the two methods. Environmental DNA detected community data exhibiting stronger associations with abiotic variables, whereas the BRUVs detected community associated more with the biotic variables.

**Fig 5.**
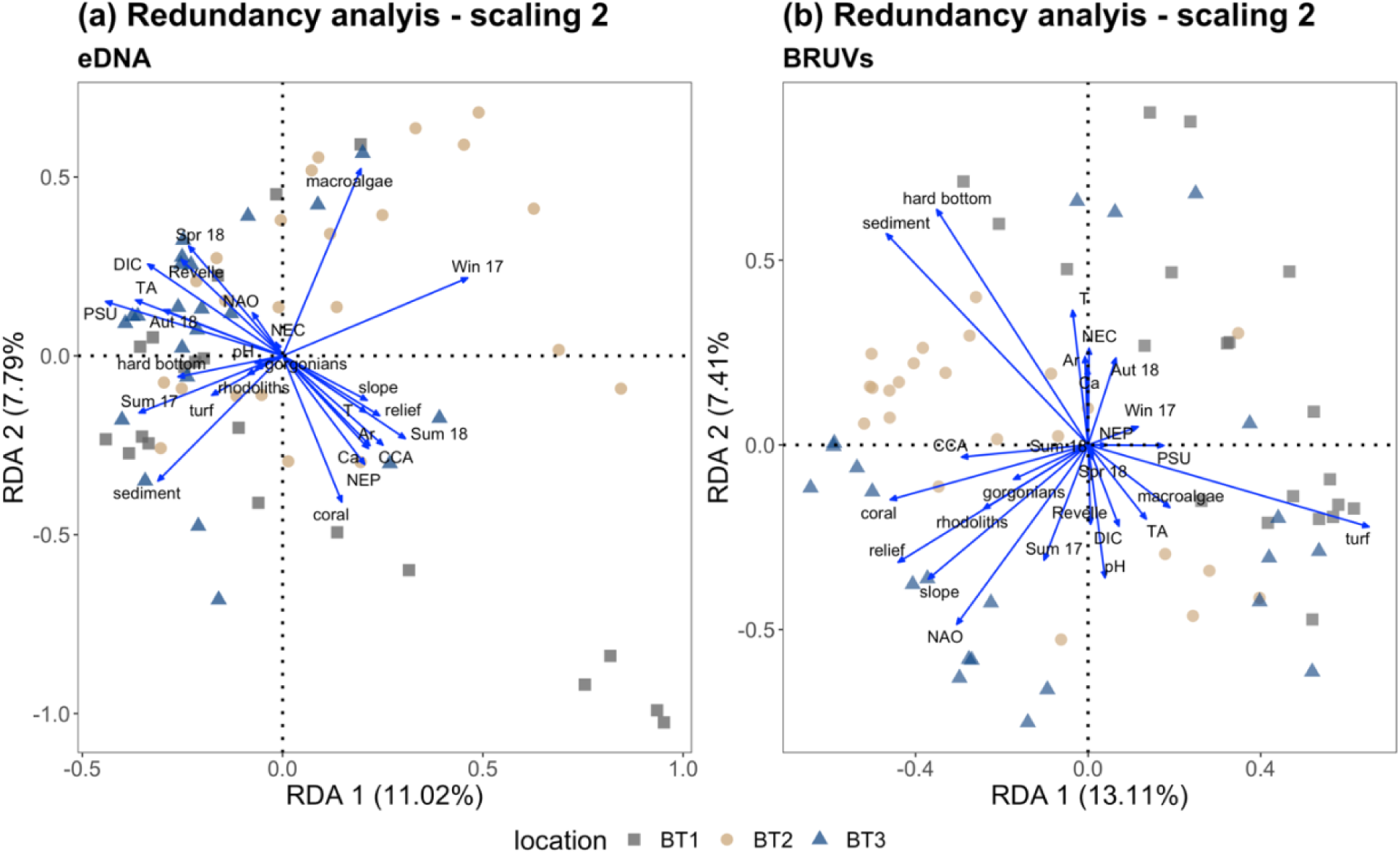
Redundancy analysis ordination diagram (triplots): location BT1 (dark grey square), BT2 (peril circles), BT3 (blue triangles). (a) environmental DNA (eDNA) metabarcoding community detection, (b) Baited Remote Underwater Video systems (BRUVs) community detection, explanatory variables (benthic composition and standardised physicochemical variables) blue arrows. First axis is horizontal, second axis is vertical. The angles among arrows denote the degree of correlation between the individual variables, and the smaller the angle, the greater the positive correlation. Negatively correlated variables are pointing in opposite directions.

All of the mesopelagic and bathypelagic fishes detected (n = 15) were only detected by the eDNA metabarcoding approach (Table S2). Of these species, half are representatives of the highly migratory lanternfish family Myctophidae. This study also recorded 38 commercially important species between the two methodologies (Table S3). The species were broadly categorised as reef dwellers (n = 25), pelagic dwellers (n=7) and baitfish (n=6). Environmental DNA detected various species of conservation interest including blue marlin, *Makaira nigricans*, a species listed as Threatened by the International Union for the Conservation of Nature Red list (IUCN; https://www.iucnredlist.org) and included in the International Commission for the Conservation of Atlantic Tunas (ICCAT). The endemic Bermuda anchovy (*Anchoa choerostoma*) was additionally detected which is listed as Endangered. BRUVs and eDNA methodologies recorded the Near Threatened lane snapper (*Lutjanus synagris*) and black grouper (*Mycteroperca bonaci*) alongside globally Vulnerable species such as grey triggerfish (*Balistes capriscus*), hogfish (*Lachnolaimus maximus*), northern red snapper (*Lutjanus campechanus*) and yellowmouth grouper (*Mycteroperca interstitialis*). Although not a fishery target species, eDNA also detected tiger shark (*Galeocerdo cuvier*), a Near Threatened highly migratory shark species, and the Critically Endangered European eel (*Anguilla anguilla*).

## Discussion

This study has demonstrated increased capacity of sampling effort through unique species detections by both methods and as a result, an additional 90 species are proposed to utilise mesophotic coral ecosystems, increasing previous research findings of Bermuda’s mesophotic ichthyofauna by 34% (Pinheiro et al. 2016; Goodbody-Gringley et al. 2019b, 2023; Stefanoudis et al. 2019).

A failure to establish complementary ecosystem-based approaches for the protection of fish and invertebrate species as stated by Strategic Goal B, Target 6 of the Aichi Biodiversity Targets, serves as a reminder of the incomplete knowledge on global biodiversity and the enormity of the challenge to rapidly obtain these data. The increased application of eDNA-based assessments over the last decade (Beng and Corlett 2020; Joydas et al. 2024) has greatly boosted the capacity to cast a much larger net as to the biodiversity present in any given ecosystem (Ruppert et al. 2019; Djurhuus et al. 2020). To date, the most comprehensive fish-centric Bermudan mesophotic studies (Pinheiro et al. 2016; Goodbody-Gringley et al. 2019b, 2023; Stefanoudis et al. 2019) have cumulatively documented 170 species ranging from 30 m into the rariphotic zone (> 150 m). These studies represent a greater collective survey effort (> 155 survey events) than this current study; yet, the combined species detections from this study totalled 182 species of which 86 where unique to this study for mesophotic focused investigations (Table S1).

Previous studies have demonstrated differences between methodologies through a greater affinity for certain taxa (Boussarie et al., 2018; Stat et al., 2019; Aglieri et al., 2021), with a tendency for eDNA biodiversity assessments to have a higher taxa detection rate over other methods likely due to the persistence of target DNA within the study area (Thomsen et al. 2012; Collins et al. 2018). The results of this study showed eDNA-derived community data exhibiting stronger associations with abiotic variables, whereas the BRUVs detected community associated more with the biotic variables. The different associations would be anticipated since eDNA samples are from a fluid environment that contains a “soup” of eDNA material from the surrounding area whereas visual methodologies (i.e., BRUVs) are strictly linked to observations that in this case, would be biased towards benthic associated species.

Whilst there was commonality of taxa across the two methods, both contributed unique detections to the overall dataset (Fig. 3; Table S1). This highlights the complementary nature of the two methodologies thus allowing for a greater accuracy of biodiversity assessment; the lack of bait species sequences in the eDNA data also shows that, with appropriate protocols in place, contamination can be avoided, and the complementary approach can be carried out by the same team.

This study revealed that habitat-driven species turnover between locations were major drivers of the fish community present at the upper/lower mesophotic interface. At each location, community diversity was dominated by species (∼ 80%) known to occur throughout the upper mesophotic and shallow reef system (Smith-Vaniz and Collette 2013). The contribution of more strictly mesophotic species accounted for ∼6% at each location. These findings suggest a high level of species continuity with the adjacent shallower reef communities despite the influence of species turnover detected between locations (Tables 2, 3). These results agree with previous findings on Bermuda mesophotic fish communities (Pinheiro et al. 2016), that documented a higher degree of turnover between 70 – 90 m, i.e., below the 60 m sites surveyed during this study. Despite the species overlap between shallow reefs and the upper mesophotic, there is increasing evidence that particular species in the Bermudan ichthyofauna are depth specialists. Bermuda chromis (*Chromis bermudae*), an endemic species, was identified using both methodologies at the upper / lower mesophotic interface, in accordance with previous studies (Stefanoudis et al., 2019; Goodbody-Gringley et al., 2019). The observations of sunshinefish (*Chromis insolata*), cherubfish (*Centropyge argi*), and tattler bass (*Serranus phoebe*) were comparable to the findings of Stefanoudis et al., (2019). This study introduces two additional specialist species longsnout butterflyfish (*Prognathodes aculeatus*) and orangeback bass (*Serranus annularis*) to the Bermudan upper / lower mesophotic interface, with the caveat that further exploration of deeper depth ranges into the rariphotic needs to be conducted. It should be noted that both these species were only detected using BRUVs observations (Table S1).

Lionfish (both species known in the area were combined; see Methods) were the only species detected in all eDNA samples suggesting a ubiquitous distribution across the 60 m depth range. However, it was only observed on 37 of the 67 BRUVs deployments. Based on behavioural observations of lionfish made during the annotation of video footage, it is plausible that individuals are present but out of the field of view. Lionfish are known to be crepuscular predators in both their native and invaded ranges (Cure et al. 2012). Given the known benefits of eDNA metabarcoding applications for detecting cryptic species (West et al. 2020; Bessey et al. 2023), the potential detection bias of lionfish would be expected. Of those species only detected by eDNA, 31% exhibit nocturnal behaviour.

Two other abundant species, – bluehead wrasse (*Thalassoma bifasciatum*) and mackerel scad (*Decapterus macarellus*), – were detected in ∼90% of the eDNA samples (100% at genus level). The bluehead wrasse is a facultative cleaner species that provides valuable ecosystem functions and was determined to be the top teleost prey species of lionfish (Eddy et al. 2016). Mackerel scad are targeted by fishers as preferred bait for demersal fishing in mesophotic ecosystems, albeit not on an industrial scale. The flyingfish genus *Cheilopogon* accounted for ∼ 50% of the eDNA detections at genus level. Flyingfish were not observed by BRUVs but their known depth ranges suggest this species is generally close to the sea surface (< 20 m). A likely explanation for the higher read count is the predator avoidance behaviour employed by this genus. As the common name suggests, flyingfish use modified pectoral fins to “fly” away from predators before collapsing their wings and splashing back into the water. This action is very likely to sluff off eDNA material and, in time, fall as “marine snow” into the 60 m depth strata being surveyed by this study. Fyingfish are ubiquitous within the surface waters above Bermudan mesophotic reefs and the surrounding pelagic environment, advection of eDNA from this genus from surface to deep would be likely.

Elasmobranchs had a noticeably low detection rate within the two datasets with each method detecting one species, Galapagos shark (*Carcharhinus galapagensis*; BRUVs) and tiger shark (*Galeocerdo cuvier*; eDNA). The known lack of affinity of the chosen MiFish-U primers for amplifying elasmobranchs (Miya et al. 2015), meant low detections through eDNA were not a surprise. Since the commencement of this study, the less-than-ideal affinity for elasmobranchs has been addressed through the development of the Elas02 primers (Taberlet et al. 2018) which are modified MiFish primers. The Elas02 primer set’s higher affinity for elasmobranchs were demonstrated in a recent assessment of shark and ray biodiversity of Reunion Island (Mariani et al. 2021a). The single visual observation of the Galapagos shark was unexpected. Baited camera (BRUVs) methodology has been adopted by Global FinPrint (Simpfendorfer et al., 2023) due to the affinity for sharks to be attracted to the bait. One explanation for the lack of observations could be due to behaviour patterns of reef sharks. Papastamatiou et al., (2015a) observed a reverse diel pattern in Galapagos shark behaviour with individuals migrating to the shallow reefs systems during the day and returning to mesophotic reefs during dark periods. All BRUVs deployments took place exclusively during the day to utilize ambient light at depth. Therefore, the combination of animal movement away from the target depth strata and low specificity of eDNA would make detections for either method less than optimal.

The ∼ 60 m depth range has historically been an important area for the local commercial fishing industry and a target area for the deployment of fish pots. In 1990, the Government of Bermuda proactively banned the use of fish pots (Butler et al. 1993) in an effort to protect reef fish species that had either demonstrated sharp declines in abundances (e.g., groupers) or increased fishing pressure (e.g., parrotfishes). As a result of this change in legislation, all parrotfishes (fam. Scaridae) where fully protected in 1993. This study detected 10 of the 14 parrotfishes accepted as part of the Bermuda ichthyofauna (Smith-Vaniz et al. 1999; Smith-Vaniz and Collette 2013), eight of which were common to both methods and one species that was unique to each method. In addition to the detection of protected species, the complementarity of these methodologies allowed for data to be gathered on the presence of commercially important species of concern (Table S3). Although the island has been designated as having advanced capacity for targeting specific species, Bermuda’s fishery is primarily for local consumption and classified as artisanal (FAO, 2022 https://www.fao.org/fishery/en/facp/bmu). However, should the island’s fishing capacity increase, the type of data presented here are likely to play a key role in enhancing future fisheries best management practices.

The detection of mesopelagic species (e.g., lanternfishes) provides a notable example of the differences in detection abilities between the two methodologies. As expected, these deeper-sea species were only detected by the molecular based approach (eDNA) and not observed by BRUVs (Table S2). It should be recognised that the inclusion of mesopelagic species in the eDNA dataset is not an admission that these species utilise mesophotic ecosystems, rather that the DNA signatures were present at the time of sampling. This is likely a direct result of the increased detection ability of eDNA over other methods and site proximity to the open ocean (∼ 1 km). However, they were retained for two reasons, 1) mesopelagic species are known to exhibit diel migration patterns, moving into the upper 200 m at night time (Dypvik and Kaartvedt 2013), a phenomenon that has been captured by eDNA metabarcoding (Canals et al. 2021), 2) these detections serve as additional biodiversity information and therefore fit with the recent analogy of Mariani et al., (2021b) on the concept of ‘molecular by-catch’.

Interestingly and unexpectedly, most moray eel species were detected by eDNA and not BRUVs. A previous assessment of Bermudan mesophotic reefs utilising BRUVs and underwater visual census (UVC), morays were only detected by BRUVs (Goodbody-Gringley et al. 2019b). It could be anticipated that the presence of bait would act as an attractant for these species since bait presence has been determined to be biased towards carnivorous fishes (Stobart et al. 2007; Lowry et al. 2012). One possible explanation could be that morays often wait to after dark to leave their core area (Kendall et al. 2021). This could be to reduce the risk of predation or due to them being primarily nocturnal predators. Clementi et al. (2021) determined a reduction in mean moray MaxN when at least one shark was also observed in the same BRUVs deployments. The use of alternative eDNA assays (e.g., Elas02; Taberlet et al. 2018) may increase elasmobranch detection and allow this to be tested in the Bermuda’s marine environment. A final noteworthy distinction between the unique species detected by each method, of the 34 shallow water reef species detected, eDNA detected all but one species (*Chaetodon ocellatus*). Baited cameras observed four unique species highlighting the complementary benefits of utilising both methodologies (Table S1).

The combined use of eDNA and BRUVs assessments allows for an increased detection of target taxa (Stat et al. 2019; Aglieri et al. 2020). In addition, the imagery generated by the BRUVs provided a secondary data stream that cannot be provided by eDNA, namely a way to quantify benthic habitat and provide a visual record of the status of that habitat. The importance of this has been demonstrated by the redundancy analysis of both the eDNA and BRUVs derived fish community data. In both cases, the explanatory variables accounted for ∼ 50% of the variability within the respective fish community datasets. This study notes there were differing community associations between the fish community datasets and benthic predictors that were likely a facet of how the fish community datasets were obtained: visual versus metabarcoding approach. Conceptually, eDNA has been subsampled on a 3D scale, i.e., water sampler (Niskin) is open top and bottom allowing water entrance from all sides. BRUVs on the other hand, is 2D since the field of view is a fixed orientation for the duration of the survey. Therefore, taxa are observed within the context of the benthic habitat and will be mostly biased by species that swim into the field of view. Environmental DNA associations are likely to reflect a broader habitat association due to either the active and / or passive transport of extracellular DNA. Irrespective of the resolution of these associations, the combination of these datasets can enhance spatial ecology and conservation modelling capacity. A point that was highlighted by Aglieri et al. (2020) through the use of eDNA to simultaneously derive functional diversity on multiple habitats and expand the power and reach of observational data. Ultimately, this would increase the effectiveness of marine spatial planning initiatives.

## Conclusion

The biomonitoring assessment of the upper/lower mesophotic interface with complementary methodologies has enabled a high-resolution evaluation of a depth zone previously established as a faunal break for mesophotic communities. The application of eDNA metabarcoding and BRUVs has enabled the detection of additional species to be considered as part of the community within this understudied ecosystem. Environmental DNA increased species and genus detection rate by 55% and 39% respectively by capturing a fluid and interactive 3-D water column. A facet that has not been included in recent publications that utilised both methodologies are the additional applications of these data that are not directly related to science. Whilst broader impacts are a ubiquitous requirement for research grants, they are often secondary to the science that is being conducted. BRUVs allow for multimedia friendly outreach material without additional costs or activities to the project and can result in novel behavioural observations that is of interest to multiple groups (Barley et al. 2016). This study has ultimately demonstrated increased capacity of sampling effort through a multi-method approach for capturing the diversity of ichthyofaunal assemblages in mesophotic coral ecosystems and increasing the number of species previously known to utilise these ecosystems.

## Acknowledgements

With the exception of the comparative data utilized from the Bermuda Atlantic Time-series Study, all data presented in this manuscript is the product of Timothy Noyes PhD thesis ‘Determining the spatial and temporal trends of mesophotic fish biodiversity and reef-scale calcification using novel approaches’, University of Salford, Manchester, UK. The following are thanked for their contribution to sampling efforts, Rosie Dowell, Jonas Schroder, Emma O’Donnell, Ellie Corbet and Alex Lundberg. Kaitlin Noyes and Christopher Noyes provided comments that improved an earlier version of this manuscript. We also thank the reviewers for constructive feedback.

## Statements and Declarations

### Author contributions

TJN: conceptualization, funding acquisition, methodology, formal analysis, visualisation, writing—original draft. LBB: conceptualization, funding acquisition, methodology, writing—review and editing, supervision. SM: writing—review and editing, supervision. GGG: conceptualization, funding acquisition, writing—review and editing. ADM: writing—review and editing, supervision. All authors gave the final approval for the publication.

### Data availability

The datasets used and/or analysed during the current study are available from the corresponding author upon reasonable request.

### Ethics approval

The study was carried out according to ethical guidelines stated by the Government of Bermuda with samples collected under Special Permit SP190603.

## References

Aglieri G, Baillie C, Mariani S, et al (2020) Environmental DNA effectively captures functional diversity of coastal fish communities. Mol Ecol 1–13. 10.1111/mec.15661

Albins MA, Hixon MA (2008) Invasive Indo-Pacific lionfish Pterois volitans reduce recruitment of Atlantic coral-reef fishes. Mar Ecol Prog Ser 367:233–238. 10.3354/meps07620

Baldwin CC, Tornabene L, Robertson DR (2018) Below the Mesophotic. Sci Rep 8:. 10.1038/s41598-018-23067-1

Barley SC, Mehta RS, Meeuwig JJ, Meekan MG (2016) To knot or not? Novel feeding behaviours in moray eels. Marine Biodiversity 46:703–705. 10.1007/s12526-015-0404-y

Baselga A, Orme CDL (2012) Betapart: An R package for the study of beta diversity. Methods Ecol Evol 3:808–812. 10.1111/j.2041-210X.2012.00224.x

Bejarano I, Appeldoorn RS, Nemeth M (2014) Fishes associated with mesophotic coral ecosystems in La Parguera, Puerto Rico. Coral Reefs 33:313–328. 10.1007/s00338-014-1125-6

Beng KC, Corlett RT (2020) Applications of environmental DNA (eDNA) in ecology and conservation: opportunities, challenges and prospects. Biodivers Conserv 29:2089–2121

Bessey C, Depczynski M, Goetze JS, et al (2023) Cryptic biodiversity: A portfolio-approach to coral reef fish surveys. Limnol Oceanogr Methods 21:594–605. 10.1002/lom3.10567

Blanco-Bercial L (2020) Metabarcoding Analyses and Seasonality of the Zooplankton Community at BATS. Front Mar Sci 7:1–16. 10.3389/fmars.2020.00173

Bohmann K, Evans A, Gilbert MTP, et al (2014) Environmental DNA for wildlife biology and biodiversity monitoring. Trends Ecol Evol 29:358–367. 10.1016/j.tree.2014.04.003

Bokulich NA, Subramanian S, Faith JJ, et al (2013) Quality-filtering vastly improves diversity estimates from Illumina amplicon sequencing. Nat Methods 10:57–59. 10.1038/nmeth.2276

Boussarie G, Bakker J, Wangensteen OS, et al (2018) Environmental DNA illuminates the dark diversity of sharks. Sci Adv 4:. 10.1126/sciadv.aap9661

Butler JN, Bumett-Herkes J, Barnes JA, Ward J (1993) The bermuda fisheries a tragedy of the commons averted? Environment 35:7–33. 10.1080/00139157.1993.9929067

Canals O, Mendibil II, Santos MM, et al (2021) Vertical stratification of environmental DNA in the open ocean captures ecological patterns and behavior of deep-sea fishes. 6. 10.1101/2021.02.10.430594

Clementi GM, Bakker J, Flowers KI, et al (2021) Moray eels are more common on coral reefs subject to higher human pressure in the greater Caribbean. iScience 24:. 10.1016/j.isci.2021.102097

Collins RA, Wangensteen OS, O’Gorman EJ, et al (2018) Persistence of environmental DNA in marine systems. Commun Biol 1:185. 10.1038/s42003-018-0192-6

Cure K, Benkwitt CE, Kindinger TL, et al (2012) Comparative behavior of red lionfish Pterois volitans on native Pacific versus invaded Atlantic coral reefs. Mar Ecol Prog Ser 467:181–192. 10.3354/meps09942

Deiner K, Bik HM, Mächler E, et al (2017) Environmental DNA metabarcoding: Transforming how we survey animal and plant communities. Mol Ecol 26:5872–5895

Djurhuus A, Closek CJ, Kelly RP, et al (2020) Environmental DNA reveals seasonal shifts and potential interactions in a marine community. Nat Commun 11:1–9. 10.1038/s41467-019-14105-1

Dorman SR, Harvey ES, Newman SJ (2012) Bait effects in sampling coral reef fish assemblages with stereo-BRUVs. PLoS One 7:1–12. 10.1371/journal.pone.0041538

Dray AS, Bauman D, Blanchet G, et al (2021) Package ‘ adespatial.’ 10.1890/11-1183.1>.Maintainer

Dypvik E, Kaartvedt S (2013) Vertical migration and diel feeding periodicity of the skinnycheek lanternfish (Benthosema pterotum) in the Red Sea. Deep Sea Res 1 Oceanogr Res Pap 72:9–16. 10.1016/j.dsr.2012.10.012

Eddy C, Pitt J, Morris JA, et al (2016) Diet of invasive lionfish (Pterois volitans and P. miles) in Bermuda. Mar Ecol Prog Ser 558:193–206. 10.3354/meps11838

Faiella G (2003) Fishing in Bermuda. Macmillian Education, Oxford

Froese R, Pauly D (2022) FishBase. In: World Wide Web electronic publication. www.fishbase.org. Accessed 20 Mar 2004

Fukunaga A, Kosaki RK, Wagner D, Kane C (2016) Structure of mesophotic reef fish assemblages in the Northwestern Hawaiian Islands. PLoS One 11:. 10.1371/journal.pone.0157861

Gold Z, Sprague J, Kushner DJ, et al (2020) eDNA metabarcoding as a biomonitoring tool for marine protected areas. bioRxiv preprint 258889:1–34. 10.1101/2020.08.20.258889

Goldberg CS, Turner CR, Deiner K, et al (2016) Critical considerations for the application of environmental DNA methods to detect aquatic species. Methods Ecol Evol 7:1299–1307. 10.1111/2041-210X.12595

Goodbody-Gringley G, Chequer A, Grincavitch C, et al (2023) Impacts of recurrent culling of invasive lionfish on mesophotic reefs in Bermuda. Coral Reefs 42:443–452. 10.1007/s00338-023-02354-y

Goodbody-Gringley G, Eddy C, Pitt JM, et al (2019a) Ecological Drivers of Invasive Lionfish (Pterois volitans and Pterois miles) Distribution Across Mesophotic Reefs in Bermuda. Front Mar Sci 6:1–12. 10.3389/fmars.2019.00258

Goodbody-Gringley G, Noyes T, Smith SR (2019b) Bermuda. In: Yossi L, Puglise KA, Bridge T (eds) Mesophotic Coral Ecosystems, Coral Reefs of the World 12. Springer, Cham, pp 31–45

Harvey ES, Cappo M, Butler JJ, et al (2007) Bait attraction affects the performance of remote underwater video stations in assessment of demersal fish community structure. Mar Ecol Prog Ser 350:245–254. 10.3354/meps07192

Hinderstein LM, Marr JCA, Martinez FA, et al (2010) Theme section on “Mesophotic Coral Ecosystems: Characterization, Ecology, and Management.” Coral Reefs 29:247–251

Holmlund CM, Hammer M (1999) Ecological Economics Ecosystem services generated by fish populations. Ecological Economics 29:253–268

Hsieh TC, Ma KH, Chao A (2016) iNEXT: an R package for rarefaction and extrapolation of species diversity (Hill numbers). Methods Ecol Evol 7:1451–1456. 10.1111/2041-210X.12613

Joydas T V., Manokaran S, Gopi J, et al (2024) Advancing ecological assessment of the Arabian Gulf through eDNA metabarcoding: opportunities, prospects, and challenges. Front Mar Sci 11

Juhel JB, Utama RS, Marques V, et al (2020) Accumulation curves of environmental DNA sequences predict coastal fish diversity in the coral triangle: EDNA predict fish diversity. Proceedings of the Royal Society B: Biological Sciences 287. 10.1098/rspb.2020.0248rspb20200248

Kelly RP, Closek CJ, O’Donnell JL, et al (2017) Genetic and manual survey methods yield different and complementary views of an ecosystem. Front Mar Sci 3:1–11. 10.3389/FMARS.2016.00283

Kelly RP, Port JA, Yamahara KM, Crowder LB (2014) Using environmental DNA to census marine fishes in a large mesocosm. PLoS One 9. 10.1371/journal.pone.0086175

Kendall MS, Siceloff L, Ruffo A, et al (2021) Green morays (Gymnothorax funebris) have sedentary ways in mangrove bays, but also ontogenetic forays to reef enclaves. Environ Biol Fishes 104:. 10.1007/s10641-021-01137-0

Legendre P, Gallagher ED (2001) Ecologically meaningful transformations for ordination of species data. Oecologia 129:271–280. 10.1007/s004420100716

Lesser MP, Slattery M, Laverick JH, et al (2019) Global community breaks at 60 m on mesophotic coral reefs. Global Ecology and Biogeography 28:1403–1416. 10.1111/geb.12940

Logan A (1988) The Holocene Reefs of Bermuda. Comparative Sedimentology Laboratory, Miami Beach (Fla.)

Logan A, Murdoch T (2011) Bermuda. In: Hopley D (ed) Encyclopedia of modern coral reefs: structure, form and process, Earth Science Series., 1st edn. Springer-Verlag, Dordrecht, pp 469–486

Lowry M, Folpp H, Gregson M, Suthers I (2012) Comparison of baited remote underwater video (BRUV) and underwater visual census (UVC) for assessment of artificial reefs in estuaries. J Exp Mar Biol Ecol 416– 417:243–253. 10.1016/j.jembe.2012.01.013

Loya Y, Puglise KA, Bridge TCL (eds) (2019) Mesophotic Coral Ecosystems. Springer International Publishing, Cham

Mächler E, Osathanunkul M, Altermatt F (2018) Shedding light on eDNA: neither natural levels of UV radiation nor the presence of a filter feeder affect eDNA-based detection of aquatic organisms. PLoS One 13:. 10.1371/journal.pone.0195529

Mariani S, Fernandez C, Baillie C, et al (2021a) Shark and ray diversity, abundance and temporal variation around an Indian Ocean Island, inferred by eDNA metabarcoding. Conserv Sci Pract 3:1–10. 10.1111/csp2.407

Mariani S, Harper LR, Collins RA, et al (2021b) Estuarine molecular bycatch as a landscape-wide biomonitoring tool. Biol Conserv 261:109287. 10.1016/j.biocon.2021.109287

Miya M, Gotoh RO, Sado T (2020) MiFish metabarcoding: a high-throughput approach for simultaneous detection of multiple fish species from environmental DNA and other samples. Springer Japan

Miya M, Sato Y, Fukunaga T, et al (2015) MiFish, a set of universal PCR primers for metabarcoding environmental DNA from fishes: detection of more than 230 subtropical marine species. R Soc Open Sci 2:150088. 10.1098/rsos.150088

Morris J a, Green SJ (2012) Lionfish Research: Current Findings an Remaining Questions

Ogle DH, Doll JC, Wheeler P, A D (2021) FSA: Fisheries Stock Analysis. R package version 0.9.1

Oksanen AJ, Blanchet FG, Friendly M, et al (2019) Vegan. Encyclopedia of Food and Agricultural Ethics 2395– 2396. 10.1007/978-94-024-1179-9_301576

Page HN, Andersson AJ, Jokiel PL, et al (2016) Differential modification of seawater carbonate chemistry by major coral reef benthic communities. Coral Reefs 35:1311–1325. 10.1007/s00338-016-1490-4

Papastamatiou YP, Meyer CG, Kosaki RK, et al (2015) Movements and foraging of predators associated with mesophotic coral reefs and their potential for linking ecological habitats. Mar Ecol Prog Ser 521:155–170. 10.3354/meps11110

Pinheiro HT, Goodbody-Gringley G, Jessup ME, et al (2016) Upper and lower mesophotic coral reef fish communities evaluated by underwater visual censuses in two Caribbean locations. Coral Reefs 35:139–151. 10.1007/s00338-015-1381-0

Port JA, O’Donnell JL, Romero-Maraccini OC, et al (2016) Assessing vertebrate biodiversity in a kelp forest ecosystem using environmental DNA. Mol Ecol 25:527–541. 10.1111/mec.13481

Porter TM, Hajibabaei M (2018) Over 2.5 million COI sequences in GenBank and growing. PLoS One 13:1–16. 10.1371/journal.pone.0200177

Pyle RL, Boland R, Bolick H, et al (2016) A comprehensive investigation of mesophotic coral ecosystems in the Hawaiian Archipelago. PeerJ 4:e2475. 10.7717/peerj.2475

R Core Team (2019) R: A language and environment for statistical computing

Rognes T, Flouri T, Nichols B, et al (2016) VSEARCH: A versatile open source tool for metagenomics. PeerJ 2016:1–22. 10.7717/peerj.2584

Rosa MR, Alves AC, Medeiros DV, et al (2016) Mesophotic reef fish assemblages of the remote St. Peter and St. Paul’s Archipelago, Mid-Atlantic Ridge, Brazil. Coral Reefs 35:113–123. 10.1007/s00338-015-1368-x

Ruppert KM, Kline RJ, Rahman MS (2019) Past, present, and future perspectives of environmental DNA (eDNA) metabarcoding: A systematic review in methods, monitoring, and applications of global eDNA. Glob Ecol Conserv 17:e00547. 10.1016/j.gecco.2019.e00547

Santana-Garcon J, Newman SJ, Harvey ES (2014) Development and validation of a mid-water baited stereo-video technique for investigating pelagic fish assemblages. J Exp Mar Biol Ecol 452:82–90. 10.1016/j.jembe.2013.12.009

Sato Y, Miya M, Fukunaga T, et al (2018) MitoFish and mifish pipeline: A mitochondrial genome database of fish with an analysis pipeline for environmental DNA metabarcoding. Mol Biol Evol 35:1553–1555. 10.1093/molbev/msy074

Schloss PD, Westcott SL, Ryabin T, et al (2009) Introducing mothur: Open-source, platform-independent, community-supported software for describing and comparing microbial communities. Appl Environ Microbiol 75:7537–7541. 10.1128/AEM.01541-09

Schobernd ZH, Bacheler NM, Conn PB (2014) Examining the utility of alternative video monitoring metrics for indexing reef fish abundance. Canadian Journal of Fisheries and Aquatic Sciences 71:464–471. 10.1139/cjfas-2013-0086

Smith-Vaniz W, Collette B (2013) Fishes of Bermuda. Smith-Vaniz, William and Collette, Bruce B 19:165

Smith-Vaniz WF, Collette BB, Luckhurst BE (1999) Fishes of Bermuda: history, zoogeography,annotated checklist, and identification keys. American Society of Ichthyologists and Herpetologists

Stat M, Huggett MJ, Bernasconi R, et al (2017) Ecosystem biomonitoring with eDNA: Metabarcoding across the tree of life in a tropical marine environment. Sci Rep 7:1–11. 10.1038/s41598-017-12501-5

Stat M, John J, DiBattista JD, et al (2019) Combined use of eDNA metabarcoding and video surveillance for the assessment of fish biodiversity. Conservation Biology 33:196–205. 10.1111/cobi.13183

Stefanoudis P V., Gress E, Pitt JM, et al (2019) Depth-dependent structuring of reef fish assemblages from the shallows to the rariphotic zone. Front Mar Sci 6:1–16. 10.3389/fmars.2019.00307

Stobart B, García-Charton JA, Espejo C, et al (2007) A baited underwater video technique to assess shallow-water Mediterranean fish assemblages: Methodological evaluation. J Exp Mar Biol Ecol 345:158–174. 10.1016/j.jembe.2007.02.009

Strickler KM, Fremier AK, Goldberg CS (2015) Quantifying effects of UV-B, temperature, and pH on eDNA degradation in aquatic microcosms. Biol Conserv 183:85–92. 10.1016/j.biocon.2014.11.038

Taberlet P, Bonin A, Zinger L, Coissac E (2018) Environmental DNA: For Biodiversity Research and Monitoring. Oxford University PressOxford, UK

Takahara T, Minamoto T, Yamanaka H, et al (2012) Estimation of fish biomass using environmental DNA. PLoS One 7:3–10. 10.1371/journal.pone.0035868

Thomsen P, Kielgast J, Iversen L, et al (2012) Detection of a Diverse Marine Fish Fauna Using Environmental DNA from Seawater Samples. PLoS One 7:1–9. 10.1371/journal.pone.0041732

Weltz K, Lyle JM, Ovenden J, et al (2017) Application of environmental DNA to detect an endangered marine skate species in the wild. PLoS One 12:1–16. 10.1371/journal.pone.0178124

West KM, Stat M, Harvey ES, et al (2020) eDNA metabarcoding survey reveals fine-scale coral reef community variation across a remote, tropical island ecosystem. Mol Ecol 29:1069–1086. 10.1111/mec.15382

Whitmarsh SK, Fairweather PG, Huveneers C (2017) What is Big BRUVver up to? Methods and uses of baited underwater video. Rev Fish Biol Fish 27:53–73

Wilson SK, Graham NAJ, Polunin NVC (2007) Appraisal of visual assessments of habitat complexity and benthic composition on coral reefs. Mar Biol 151:1069–1076. 10.1007/s00227-006-0538-3

Yamamoto S, Masuda R, Sato Y, et al (2017) Environmental DNA metabarcoding reveals local fish communities in a species-rich coastal sea. Sci Rep 7:. 10.1038/srep40368

Yao M, Zhang S, Lu Q, et al (2022) Fishing for fish environmental DNA: Ecological applications, methodological considerations, surveying designs, and ways forward. Mol Ecol 5132–5164. 10.1111/mec.16659

