## Supplemental data for "Complementary assessment of fish biodiversity across the upper/lower mesophotic interface in a subtropical coral reef using eDNA metabarcoding and baited cameras"

**Title:**

Timothy J. Noyes[1,2,3], 0000-0001-9750-9193

Leocadio Blanco-Bercial[1,2], 0000-0003-0658-7183

Stefano Mariani4, 0000-0002-5329-0553

Gretchen Goodbody-Gringley5 0000-0003-3439-0919

Allan D. McDevitt3,6 0000-0002-2677-7833

**Addresses:**

[1]Bermuda Institute of Ocean Sciences, Arizona State University (ASU), St. George's, Bermuda

[2]School of Ocean Futures, Julie Ann Wrigley Global Futures Laboratory, Arizona State University (ASU), Tempe, Arizona, USA

[3]School of Science, Engineering and Environment, University of Salford, Salford, UK

[4]School of Biological and Environmental Sciences, Liverpool John Moores University, Liverpool, UK

[5]Central Caribbean Marine Institute, Little Cayman Island, Cayman Islands, UK

[6]Marine and Freshwater Research Centre, Department of Natural Resources and the Environment, Atlantic Technological University, Galway, Ireland

***Laboratory methods***

DNA extractions from the frozen filters were performed using the E.Z.N.A. Mollusc DNA Kit (Omega Bio-Tek: Norcross, Georgia, USA) commercial kit. Modifications to the protocol included the use of 500µl ml buffer at step 2. Samples were incubated at 60°C for 4 hours and vortex at maximum for 1 minute every 30 minutes to aid lysis (step 3). After incubation, all samples were transferred to sterile 1.5 ml microcentrifuge tube prior to the addition of 500 µl chloroform:isoamyl alcohol (24:1; step 4). Samples were centrifugated for 14 minutes at 10,000 x g at step 5. Those samples that required two filters to obtain the desired sample volume of 8 L were pooled during step 12 of by passing the total sample through the same HiBind® DNA Mini Column. The elution step (27) was repeated twice using the same 50 µl 70°C DNA free water, in an attempt to increase DNA yield.

The first step PCR was performed using 1 µl of 10 µM of both the forward and reverse primers combined with sample volume that equated to 500 ng of DNA. Each reaction was filled up to 20 µl. Both positive and negative controls were run with all PCR steps to control for contamination. Positive control samples were sourced from DNA extracts of local reef fishes incorporated into a local reference database (Noyes and Blanco-Bercial unpublished). A SimpliAmp Thermal Cycler (Applied biosystems: Foster City, California, USA) was used for the PCR, with the following thermal profile an initial denaturation at 94°C for 3 minutes followed by 30 cycles of: 30 seconds at 94°C; annealing at 60°C for 30 seconds; extension at 69°C for 30 seconds and a final extension period at 69°C for 7 minutes. For confirmation of successful PCR, the product was run on 2 % agarose gel with a Quick-Load Purple Low Molecular Weight DNA Ladder (New England Biolabs: Massachusetts, USA). PCR product was diluted (1:10) with DNA free water and 10µl used as the template for the second PCR step, combined with 1µL of 10 µM forward and reverse dual Illumina index primers in a total reaction volume of 20 µl. The reaction thermal profile followed step one. Each second step PCR was completed in triplicate with reactions pooled to give a final volume ~55 µl (after 5 µl were used for gel electrophoresis). To allow for sample identification following demultiplexing, samples were amplified with a combination of unique forward and reverse eight base-pair indexes allowing for 64 unique dual-indexed combinations.

**Table S1.** Summary of this study in comparison to previous mesophotic fish biodiversity studies. B=Baited Remote Underwater Video Systems (BRUVs) E= environmental DNA (eDNA), X = pooled visual assessment methods (diver based. underwater visual surveys and underwater visual surveys from a submersible). Note, species data presented in Pinheiro et al. (2016) are included in data presented in Goodbody-Gringley et al (2019b).

- [a]Recorded as *Acanthurus bahianus/chirurgus* in Stefanoudis et al 2019
- [b]Recorded as *Acanthurus bahianus* in Pinheiro et al., 2016 and Goodbody-Gringley et al., 2019a
- [c]Recorded as *Chromis aff enchrysura* in Pinheiro et al., 2016; *Chromis cf. enchrysura* in Goodbody-Gringley et al., 2019a; *Chromis enchrysura* in Goodbody-Gringley et al., 2019b
- [d]Casual observation in Stefanoudis et al., 2019
- [e]Recorded as Kyphosus incisor/sectatrix in Stefanoudis et al., 2019
- [f]Recorded as *Seriola lalandi* in Goodbody-Gringley et al., 2023
- [g]Recorded as *Stegastes variabilis* in Goodbody-Gringley et al., 2019a

| Species | This Study | GGG et al., 2019a | GGG et al., 2019b | Stefanoudis et al., 2019 | GGG et al., 2023 |
|---|---|---|---|---|---|
| *Abudefduf saxatilis* | E | | | | |
| *Acanthocybium solandri* | | | X | | |
| *Acanthostracion polygonius* | B,E | | X | | |
| *Acanthostracion quadricornis* | B | | | | |
| *Acanthurus chirurgus* [a] | B | X | X | X | X |
| *Acanthurus coeruleus* | B,E | | X | | |
| *Acanthurus tractus* [b] | B,E | | X | X | X |
| *Ahlia egmontis* | E | | | | |
| *Alectic ciliaris* | | | | X | |
| *Alepisaurus ferox* | E | | | X | |
| *Aluterus monoceros* | E | | | X | |
| *Aluterus scriptus* | E | | | | |
| *Amblycirrhitus pinos* | | | | X | |
| *Anchoa choerostoma* | E | | | | |
| *Anguilla anguilla* | E | | | | |
| *Anoplogaster cornuta* | E | | | | |
| *Antennarius ocellatus* | | | | X | |
| *Anthias tenuis* | | | | X | |
| *Antigonia capros* | | | | X | |
| *Apogon evermanni* | | | X | | |
| *Apogon gouldi* | | X | | X | |
| *Apogon maculatus* | E | | | | |
| *Ariosoma balearicum* | E | | | | |
| *Aulopus filamentosus* | | | | X | |
| *Aulostomus maculatus* | B,E | | X | | X |
| *Aulostomus strigosus* | | | | X | |
| *Auxis rochei* | E | | | | |
| *Auxis thazard* | E | | | | |
| *Balistes capriscus* | E | | X | | |
| *Bathygobius curacao* | E | | | | |

| Species | This Study | GGG et al., 2019a | GGG et al., 2019b | Stefanoudis et al., 2019 | GGG et al., 2023 |
|---|---|---|---|---|---|
| *Bodianus pulchellus* | B,E | X | X | | X |
| *Bodianus rufus* | B,E | X | X | | X |
| *Bolinichthys nikolayi* | E | | | | |
| *Bothus lunatus* | E | | X | | |
| *Bothus robinsi* | E | | | | |
| *Brotula barbata* | | | | X | |
| *Calamus bajonado* | B | | X | | |
| *Calamus calamus* | B | | X | | |
| *Cantherhines macrocerus* | B | | X | | X |
| *Cantherhines pullus* | E | | X | | |
| *Canthidermis sufflamen* | E | | X | | X |
| *Canthigaster rostrata* | B,E | X | X | X | X |
| *Carangoides bartholomaei* | B,E | | X | | |
| *Caranx crysos* | E | | | | |
| *Caranx latus* | B,E | | X | | X |
| *Caranx lugubris* | B,E | | X | X | X |
| *Caranx ruber* | B,E | | X | | X |
| *Carcharhinus falciformis* | | | X | | |
| *Carcharhinus galapagensis* | B | | X | X | |
| *Caulolatilus bermudensis* | | | | X | |
| *Centropyge argi* | B,E | X | X | | X |
| *Cephalopholis cruentata* | B | | X | | |
| *Cephalopholis fulva* | B,E | X | X | | X |
| *Ceratoscopelus maderensis* | E | | | | |
| *Chaetodon capistratus* | B,E | | X | | |
| *Chaetodon ocellatus* | B | X | X | | X |
| *Chaetodon sedentarius* | B,E | X | X | | X |
| *Chaetodon striatus* | B,E | | | | |
| *Channomuraena vittata* | | | | X | |
| *Cheilopogon cyanopterus* | E | | | | |
| *Cheilopogon exsiliens* | E | | | | |
| *Chlopsis dentatus* | | | | X | |
| *Chloroscombrus chrysurus* | | | | X | |
| *Chromis bermudae* | B,E | X | X | | X |
| *Chromis vanbebberae* [c] | B | X | X | | |
| *Chromis cyanea* | B,E | X | | | X |
| *Chromis insolata* | B | | | X | |
| *Clepticus parrae* | B,E | | X | | |
| *Clupea harengus* | E | | | | |
| *Conger esculentus* | | | | X | |
| *Conger triporiceps* | | | X | X | |
| *Cookeolus japonicus* | | | | X | |
| *Cryptotomus roseus* | B,E | | | | |
| *Cyclothone pallida* | E | | | | |
| *Dactylopterus volitans* | B | | | X | |
| *Decapterus macarellus* | E | | | | |
| *Decapterus tabl* | | | | X | |
| *Decodon puellaris* | B | | X | | |
| *Diaphus dumerilii* | E | | | | |
| *Diaphus effulgens* | E | | | | |

| Species | This Study | GGG et al., 2019a | GGG et al., 2019b | Stefanoudis et al., 2019 | GGG et al., 2023 |
|---|---|---|---|---|---|
| *Diaphus mollis* | E | | | | |
| *Diodon holocanthus* | B,E | | X | | |
| *Diodon hystrix* | E | | X | | |
| *Diplodus bermudensis* | E | | X | | |
| *Diplospinus multistriatus* | E | | | | |
| *Echeneis naucrates* | E | | | | |
| *Elagatis bipinnulata* | B | | | | X |
| *Emblemaria atlantica* | E | | | | |
| *Enchelycore carychroa* | E | | | | |
| *Epinephelus adscensionis* | | | X | | |
| *Epinephelus drummondhayi* | | | X | | |
| *Epinephelus guttatus* | B,E | | X | | |
| *Epinephelus morio* | | | X | | |
| *Epinephelus mystacinus* | | | | X | |
| *Epinephelus niveatus* | | | | X | |
| *Epinephelus striatus* | | | | X | |
| *Etelis oculatus* | | | | X | |
| *Eucinostomus jonesii* | E | | | | |
| *Eustomias obscurus* | E | | | | |
| *Euthynnus alletteratus* | E | | | | |
| *Fistularia tabacaria* | B | | X | | |
| *Galeocerdo cuvier* | E | | X | | |
| *Gempylus serpens* | E | | | | |
| *Gephyroberyx darwinii* | | | | X | |
| *Gerres cinereus* | E | | | | |
| *Gnatholepis cauerensis* | E | | X | | |
| *Gnatholepis thompsoni* | | | X | | |
| *Gobiosoma macrodon* | B | | | | |
| *Gonichthys cocco* | E | | | | |
| *Gymnothorax funebris* | | X | X | | X |
| *Gymnothorax maderensis* [d] | | | | X | |
| *Gymnothorax miliaris* | E | | | | |
| *Gymnothorax moringa* | B | | X | | |
| *Gymnothorax polygonius* | E | | X | X | |
| *Gymnothorax vicinus* | E | | | | |
| *Haemulon aurolineatum* | B,E | | X | | |
| *Haemulon flavolineatum* | B,E | X | X | X | X |
| *Haemulon macrostomum* | B | | X | | |
| *Haemulon melanurum* | B,E | | | | |
| *Haemulon sciurus* | B,E | | X | | |
| *Halichoeres bathyphilus* | B | | X | | |
| *Halichoeres bivittatus* | E | | | | |
| *Halichoeres garnoti* | B | X | X | | X |
| *Halichoeres maculipinna* | E | | X | | |
| *Halichoeres radiatus* | B,E | X | X | | X |
| *Hemiramphus bermudensis* | E | | | | |
| *Heteroconger longissimus* | | | X | | |
| *Histrio histrio* | E | | | | |
| *Holacanthus bermudensis* | B,E | | X | | X |
| *Holacanthus ciliaris* | E | | | | X |
| *Holacanthus tricolor* | B,E | X | X | X | X |
| Holocanthus townsendi | | | | | X |

| Species | This Study | GGG et al., 2019a | GGG et al., 2019b | Stefanoudis et al., 2019 | GGG et al., 2023 |
|---|---|---|---|---|---|
| *Holocentrus adscensionis* | B | | X | | |
| *Holocentrus rufus* | | | X | | X |
| *Hoplostethus occidentalis* | | | | X | |
| *Hygophum hygomii* | E | | | | |
| *Hypoatherina harringtonensis* | E | | | | |
| *Isurus oxyrinchus* | | | | X | |
| *Jenkinsia lamprotaenia* | E | | | | |
| *Katsuwonus pelamis* | E | | | | |
| *Kyphosus bigibbus* | E | | | | |
| *Kyphosus cinerascens* | E | | | | |
| *Kyphosus sectatrix* [e] | B,E | | X | X | |
| *Kyphosus vaigiensis* | E | | | | |
| *Lachnolaimus maximus* | B,E | | X | | |
| *Lactophrys trigonus* | B | | X | | X |
| *Lactophrys triqueter* | B,E | | X | | |
| *Laemonema yarrellii* | E | | | X | |
| *Lagocephalus lagocephalus* | E | | | | |
| *Lagodon rhomboides* | E | | | | |
| *Lampadena atlantica* | E | | | | |
| *Lobotes surinamensis* | E | | | | |
| *Lutjanus analis* | | | X | | |
| *Lutjanus buccanella* | | | | X | |
| *Lutjanus campechanus* | E | | | | |
| *Lutjanus griseus* | B,E | | X | | |
| *Lutjanus synagris* | B,E | | X | | |
| *Lutjanus vivanus* | | | | X | |
| *Lythrypnus mowbrayi* | | | | X | |
| *Lythrypnus spilus* | | | | X | |
| *Makaira nigricans* | E | | | | |
| *Malacanthus plumieri* | B,E | | X | | X |
| *Megalops atlanticus* | E | | | | |
| *Melichthys niger* | E | X | X | | |
| *Microspathodon chrysurus* | E | | | | |
| *Mobula birostris* | | | | X | |
| *Monacanthus tuckeri* | B,E | | | | |
| *Moringua edwardsi* | E | | | | |
| *Mugil curema* | E | | | | |
| *Mulloidichthys martinicus* | B,E | | X | | |
| *Mustelus canis insularis* | | | | X | |
| *Mycteroperca bonaci* | B,E | | X | X | X |
| *Mycteroperca interstitalis* | B | | X | X | X |
| *Mycteroperca venenosa* | | | | X | |
| *Myripristis jacobus* | | | | X | |
| *Nannobrachium lineatum* | E | | | | |
| *Nemaclinus atelestos* | | | | X | |
| *Nemichthys curvirostris* | E | | | | |
| *Nomeus gronovii* | E | | | | |
| *Ocyurus chrysurus* | B,E | | X | | |
| *Ogilbia cayorum* | E | | | | |
| *Opisthonema oglinum* | E | | | | |
| *Ostichthys trachypoma* | | | | X | |
| *Paranthias furcifer* | B,E | X | X | X | X |

| **Species** | **This Study** | **GGG et al., 2019a** | **GGG et al., 2019b** | **Stefanoudis et al., 2019** | **GGG et al., 2023** |
|---|---|---|---|---|---|
| *Parasphyraenops atrimanus* | | | | X | |
| *Pempheris schomburgkii* | E | | | | |
| *Phaeoptyx conklini* | | | | X | |
| *Platybelone argalus* | E | | | | |
| *Plectrypops retrospinis* | B,E | | | X | |
| *Polymixia lowei* | | | | X | |
| *Polymixia nobilis* | | | | X | |
| *Polyprion americanus* | | | | X | |
| *Pontinus castor* | | | | X | |
| *Priacanthus arenatus* | | | | X | |
| *Priolepis hipoliti* | E | | | | |
| *Prionace glauca* | | | | X | |
| *Pristigenys alta* | | | | X | |
| *Pristipomoides macrophthalmus* | | | | X | |
| *Prognathodes aculeatus* | B | | X | | X |
| *Prognathodes cf. guyanensis* | | | | X | |
| *Pronotogrammus martinicensis* | | | | X | |
| *Pseudocaranx dentex* | B,E | | X | X | |
| *Pseudupeneus maculatus* | B,E | | X | X | X |
| *Pterois spp.* | B,E | | X | X | |
| *Pteroplatytrygon violacea* | | | | X | |
| *Regalecus glesne* | | | | X | |
| *Rypticus saponaceus* | B | X | X | | X |
| *Sardinella aurita* | E | | | | |
| *Sargocentron bullisi* | | | X | | |
| *Sargocentron coruscum* | E | | | | |
| *Scarus coelestinus* | B,E | | X | | X |
| *Scarus coeruleus* | | | X | | |
| *Scarus guacamaia* | B,E | | X | X | X |
| *Scarus iseri* | B,E | | | | |
| *Scarus taeniopterus* | B,E | X | X | | X |
| *Scarus vetula* | B,E | | X | | |
| *Schedophilus ovalis* | E | | | | |
| *Schedophilus velaini* | E | | | | |
| *Scomberomorus cavalla* | | | X | | |
| *Scorpaena albifimbria* | | | | X | |
| *Scorpaena isthmensis* | | | | X | |
| *Seriola dumerili* | B,E | | X | | |
| *Seriola fasciata* [f] | | | | X | X |
| *Seriola rivoliana* | B,E | | X | | X |
| *Serranus annularis* | B | | | X | |
| *Serranus phoebe* | B,E | | X | | |
| *Serranus tigrinus* | | | X | | X |
| *Serrivomer beanii* | E | | | | |
| *Sparisoma atomarium* | B | | | X | |
| *Sparisoma aurofrenatum* | B,E | X | | | X |
| *Sparisoma chrysopterum* | B,E | | X | | |
| *Sparisoma rubripinne* | E | | X | | |
| *Sparisoma viride* | B,E | | X | | X |
| *Sphoeroides spengleri* | B | X | X | | |
| *Sphyraena barracuda* | B,E | | X | | X |
| *Squalus cubensis* | | | | X | |

| Species | This Study | GGG et al., 2019a | GGG et al., 2019b | Stefanoudis et al., 2019 | GGG et al., 2023 |
|---|---|---|---|---|---|
| *Stegastes leucostictus* | E | | | | |
| *Stegastes partitus* | E | X | X | | |
| *Stegastes planifrons* | E | | | | |
| *Stegastes xanthurus* [9] | | X | | | |
| *Stephanolepis hispida* | E | | | | |
| *Syacium papillosum* | | | | X | |
| *Synodus foetens* | E | | X | | |
| *Synodus intermedius* | B,E | | | | |
| *Synodus synodus* | E | | | | |
| *Thalassoma bifasciatum* | B,E | X | X | | X |
| *Trachinotus goodei* | E | | | | |
| *Tylosurus crocodilus* | E | | | | |
| *Ulaema lefroyi* | E | | | | |
| *Uraspis secunda* | | | | X | |
| *Uroconger syringinus* | E | | | | |
| *Uropterygius macularius* | | | | X | |
| *Xanthichthys ringens* | B,E | | X | | |
| *Xyrichtys martinicensis* | E | | | | |
| *Xyrichtys splendens* | E | | | | |

**Table S2.** Summary of deep-sea species detected by eDNA metabarcoding. Depth ranges as published on fishbase.org (Froese and Pauly 2022). Note, (*) denotes the usual depth range preference of a species.

| Species | Common Name | Depth range (m) |
|---|---|---|
| *Alepisaurus ferox* | Long snouted lancetfish | 0 - 1830 |
| *Anoplogaster cornuta* | Common fangtooth | 500 - 200 (*) |
| *Bolinichthys nikolayi* | Lanternfish | 25 - 1760 |
| *Ceratoscopelus maderensis* | Madeira lantern fish | 51 - 1480 |
| *Cyclothone pallida* | Tan bristlemouth | 600 - 1800 (*) |
| *Diaphus effulgens* | Headlight fish | 0 - 6000 |
| *Diaphus mollis* | Soft lanternfish | 50 - 600 |
| *Diplospinus multistriatus* | Striped escolar | 50 - 1000 |
| *Eustomias obscurus* | Barbed dragonfish | 20 - 1900 |
| *Gempylus serpens* | Snake mackerel | 0 - 600 |
| *Gonichthys cocco* | Cocco's lanternfish | 425 - 650 (*) |
| *Hygophum hygomii* | Bermuda lanternfish | 0 - 1485 |
| *Lampadena atlantica* | Lanternfish | 60 - 1000 |
| *Nannobrachium lineatum* | Lanternfish | 60 - 1150 |
| *Nemichthys curvirostris* | Pale threadtail snipe eel | 0 - 2000 |

**Table S3.** Summary of 38 commercially important species listed by IUCN status (colour-coded according to status) and the method of detection used to record presence.

| Species | Common name | Origin | BRUVs only | eDNA only | BRUVs and eDNA | IUCN Status |
|---|---|---|---|---|---|---|
| *Anchoa choerostoma* | Bermuda anchovy | Baitfish | | X | | Endangered |
| *Auxis rochei* | Bullet tuna | Pelagic | | X | | Least Concern |
| *Auxis thazard* | Skipjack tuna | Pelagic | | X | | Least Concern |
| *Balistes capriscus* | Grey triggerfish | Reef | | X | | Vulnerable |
| *Calamus bajonado* | Jolthead porgy | Reef | X | | | Least Concern |
| *Caranx latus* | Horse-eye jack | Reef | | | X | Least Concern |
| *Caranx lugubris* | Black jack | Reef | | | X | Least Concern |
| *Caranx ruber* | Bar jack | Reef | | | X | Least Concern |
| *Carcharhinus galapagensis* | Galpagos shark | Reef | X | | | Least Concern |
| *Cephalopholis fulva* | Coney | Reef | | | X | Least Concern |
| *Clupea harengus* | Atlantic Herring | Baitfish | | X | | Least Concern |
| *Decapterus macarellus* | Mackerel scad | Baitfish | | X | | Least Concern |
| *Diplodus bermudensis* | Bermuda bream | Reef | | X | | Least Concern |
| *Elagatis bipinnulata* | Rainbow runner | Pelagic | X | | | Least Concern |
| *Epinephelus guttatus* | Red hind | Reef | | | X | Least Concern |
| *Euthynnus alletteratus* | Little tunny | Pelagic | | X | | Least Concern |
| *Haemulon flavolineatum* | French grunt | Reef | | | X | Least Concern |
| *Haemulon sciurus* | Bluesstriped grunt | Reef | | | X | Least Concern |
| *Hemiramphus bermudensis* | Bermuda halfbeak | Reef | | X | | Least Concern |
| *Hypoatherina harringtonens* | Reef silverside | Baitfish | | X | | Least Concern |
| *Jenkinsia lamprotaenia* | Dwarf round herring | Baitfish | | X | | Least Concern |
| *Katsuwonus pelamis* | Skipjack tuna | Pelagic | | X | | Least Concern |
| *Lachnolaimus maximus* | Hogfish | Reef | | | X | Vulnerable |
| *Lutjanus campechanus* | Northern red snapper | Reef | | X | | Vulnerable |
| *Lutjanus griseus* | Grey snapper | Reef | | | X | Least Concern |
| *Lutjanus synagris* | Lane snapper | Reef | | | X | Near Threatened |
| *Makaira nigricans* | Blue Marlin | Pelagic | | X | | Vulnerable |
| *Mugil curema* | White mullet | Reef | | X | | Least Concern |
| *Mycteroperca bonaci* | Black grouper | Reef | | | X | Near Threatened |
| *Mycteroperca interstitialis* | Yellowmouth grouper | Reef | X | | | Vulnerable |
| *Ocyurus chrysurus* | Yellowtail snapper | Reef | | | X | Data Deficient |
| *Opisthonema oglinum* | Atlantic thread herring | Baitfish | | X | | Least Concern |
| *Paranthias furcifer* | Creolefish/barber | Reef | | | X | Least Concern |
| *Pseudocaranx dentex* | White trevally | Reef | | | X | Least Concern |
| *Sardinella aurita* | Round sardinella | Baitfish | | X | | Least Concern |
| *Seriola dumerili* | Greater amberjack | Reef | | | X | Least Concern |
| *Seriola rivoliana* | Almaco jack/bonita | Reef | | | X | Least Concern |
| *Sphyraena barracuda* | Barracuda | Reef | | | X | Least Concern |

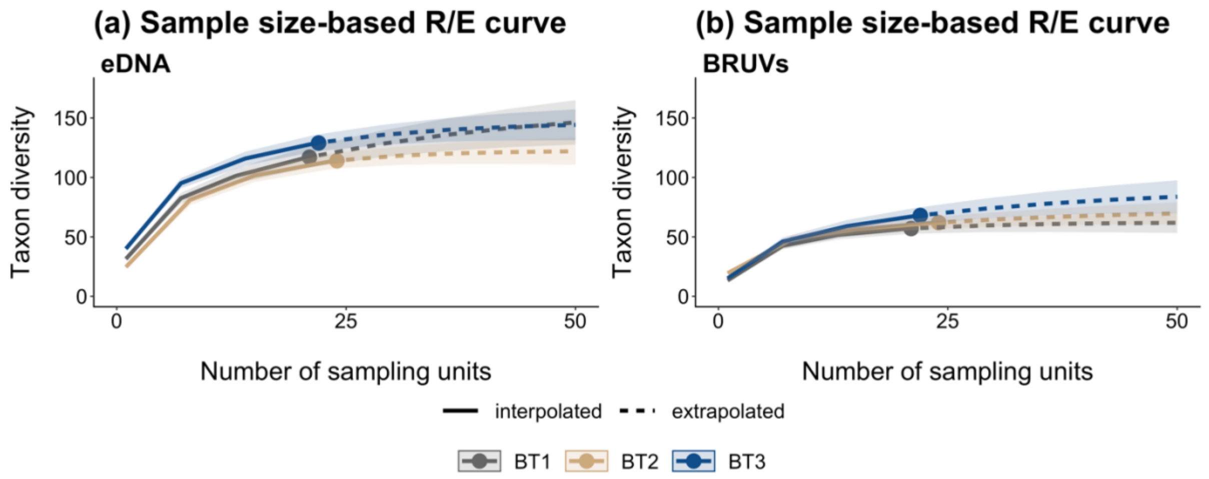


**Fig. S1.** Sample size-based rarefaction/extrapolation (R/E) curves for each location BT1 (dark gray symbols and lines), BT2 (peril symbols and lines), BT3 (blue symbols and lines): (**a**) environmental DNA (eDNA) metabarcoding methodology, (**b**) Baited Remote Underwater Video systems (BRUVs) methodology.
